# Correlated dynamics, reactive conformations and non-Arrhenius behaviour in the temperature-dependence of enzyme activity: triosephosphate isomerase

**DOI:** 10.64898/2025.12.31.695415

**Authors:** Swati Singh, Dan Pike, Erica J. Prentice, Tom A. Williams, Vickery L. Arcus, Marc W. van der Kamp, Adrian J. Mulholland

## Abstract

Heat capacity changes are increasingly recognised as crucial for understanding the temperature dependence of enzyme-catalysed reaction rates. Here, we combine experiments and molecular dynamics simulations to investigate triosephosphate isomerase (TIM, TPI) from psychrophilic, mesophilic and thermophilic organisms. Kinetic data show clear curvature in rate-temperature plots, particularly for the cold-adapted enzyme, independent of unfolding. This non-Arrhenius behaviour is accounted for by Macromolecular Rate Theory (MMRT), implying an activation heat capacity, largest for the psychrophile. Simulations reveal the molecular origins of these differences, showing significant conformational and dynamical changes between reactant and transition states in the cold-adapted enzyme, with a smaller activation heat capacity in the mesophilic enzyme, and near-zero for the thermophile. Transition-state-like conformations and reorganized correlated dynamics underlie these adaptations. These results demonstrate that dynamical differences between crucial states along the reaction pathway play a key role in determining the optimum temperature of activity for this archetypal enzyme.

## Introduction

Life has a remarkable ability to adapt to diverse habitats. One of the most striking adaptations is the range of temperatures at which life exists on Earth. The molecular origins of these evolutionary adaptations remain a major open question. Psychrophilic and thermophilic organisms grow at low or high temperatures, in which other organisms typically cannot grow and/or survive. This adaptation requires many sequence, structural and physiological adjustments (*1–3*) but central to thermal adaptation is the temperature dependence of enzyme activity (*4*). Survival at high temperatures obviously requires thermal stability; cold adaptation requires activity at low temperatures at which less thermal energy is available.

Enzymes typically have a temperature of maximum activity, the peak of which is correlated with, but generally differs from, the optimal growth conditions of the organism. The temperature of maximum activity is often higher than the optimal growth temperature of the organism, which instead coincides with an inflection point in enzyme activity-temperature curves (*5*). Above enzyme temperature optimum, their activity declines, generally *before* unfolding. Understanding this behaviour is fundamentally important in molecular evolution and for design and engineering of protein catalysts for practical applications.

Growing evidence suggests that the temperature-dependent behaviour of enzyme activity is more complex than previously thought. Activation heat capacity is becoming pivotal in understanding how enzyme-catalysed reaction rates depend on temperature. This is formalized in Macromolecular Rate Theory (MMRT) (*4*, *6–10*).

Loss of enzyme activity at high temperatures is traditionally attributed to protein unfolding (*11*). Unfolding alone does not account for observed behaviour, however: e.g., psychrophilic enzymes typically remain folded and active above their optimum temperatures but have reduced activity (*12*). Structurally, psychrophilic enzymes are generally similar to their mesophilic and thermophilic equivalents, particularly in their active sites (*13*, *14*). There are important differences in activation enthalpy and entropy linked to cold temperature adaptation (*13*, *15*) : lower activation enthalpy values coupled with compensatory changes in activation entropy. Wolfenden has described this as “enthalpic catalysis”, which is increasingly favoured as temperature decreases (*16*). Experimental and simulation studies have corroborated this relationship between activation enthalpy and entropy (*15*). Psychrophilic enzymes exhibit reduced activation enthalpies (with a trade-off of lower activation entropies) (*15*, *17–19*) Åqvist *et al.* found, for simulations of homologous citrate synthases and triosephosphate isomerases, that this is caused by changes in flexibility at the enzyme surface (*17*, *20*, *21*). Such adaptations are crucial to adaptation to different temperatures, but they do not account for curvature (non-Arrhenius behaviour) observed in experimental temperature-rate plots.

Here, we analyse triosephosphate isomerase (TIM, TPI) from thermophilic, mesophilic and psychrophilic organisms, to investigate the factors that govern the temperature dependence of activity. TIM is an exquisitely evolved, usually dimeric enzyme involved in glycolysis, interconverting dihydroxyacetone phosphate and D-glyceraldehyde-3-phosphate (*20*). Beyond its glycolytic function, TIM exhibits diverse roles, including as a virulence factor in some pathogenic organisms; a potential allergen; roles in sperm function and fertility; and is implicated in cancer progression and Alzheimer’s disease pathology (*21*). TIM is remarkably efficient and has been famously described as “catalytically perfect” (*22*, *23*): its turnover has been argued to be limited by diffusion (*22*, *24*) It gives its name to the TIM barrel (consisting of eight α-helices and eight parallel β-strands), one of the most frequently observed natural protein folds. This fold is highly versatile and evolvable, accommodating diverse catalytic functions. (*25–27*). Loop dynamics play a critical role in regulating activity and evolvability in TIM barrel enzymes (*28*). It has been argued to have facilitated the early evolution of enzyme-mediated metabolism, making it an excellent model system (*7*, *17*). Oligomerisation plays a critical role in TIM catalytic efficiency: engineered monomeric variants show substantially lower activity than the native dimeric form, though monomeric enzymes can retain basal catalytic function (*20*). This enhancement appears to involve coordinated conformational dynamics between subunits, particularly affecting the essential closure of loop 6 over the active site during catalysis (*29*, *30*). Evidence indicates that dimerization enables the collective motions necessary for optimal enzyme function, helping TIM achieve its remarkably high catalytic efficiency approaching the diffusion limit (*31*). In some hyperthermophilic organisms, TIM forms tetramers that provide additional thermal stability while maintaining the coordinated loop dynamics essential for catalytic function (*32*). Here, we analyse dimeric TIM enzymes from thermophilic (hot-TIM, from *Thermoplasma acidophilum*), mesophilic (meso-TIM, from *Saccharomyces cerevisiae*), and psychrophilic organisms (cold-TIM, from *Vibrio marinus*), to analyse their temperature-dependent activity and dynamics.

We measure activity over a wide temperature range in high resolution kinetics experiments: these show clear and significant curvature in the temperature dependence of activity for cold-and meso-TIM, independent of unfolding and not due to changes in rate-limiting step. Molecular dynamics (MD) simulations show significant differences in conformational and dynamical behaviour between transition state analogue (TSA) and reactant state (RS) complexes, identifying a “transition-state-like conformation” (TLC) (*33*). The MD simulations are also used to calculate activation heat capacities (Δ*C*^‡^), which are found to be largest in magnitude for the cold-adapted TIM, smallest for hot-TIM, and intermediate for meso-TIM intermediate. These findings are consistent with the predictions of MMRT, i.e. that catalysis at low temperatures will be accompanied by increased conformational fluctuations and increasingly negative values of Δ*C*^‡^ (*6*). The simulations here identify dynamical causes of this behaviour, showing significant differences in correlated dynamics for the TLC in cold-TIM, and also in meso-TIM (but not in hot-TIM). These combined experimental and simulation results indicate that activation heat capacity is centrally important to temperature adaptation of activity in this prototypical enzyme.

## Results

### Non-Arrhenius behaviour

High-resolution kinetics experiments on the temperature dependence of the activity of cold-TIM and meso-TIM (details in SI) show clear and significant curvature (non-Arrhenius behaviour), particularly for cold-TIM. We tested fitting of these data with MMRT and its variants (**Figure 1, Figure S1 and Table S1**). Activation heat capacity (Δ*C*^‡^) can be calculated by fitting the natural logarithm of the rate-versus-temperature plot using MMRT (*4*, *6*, *34*). For cold-TIM, the data were fit somewhat better using a linearly temperature-dependent Δ*C*^‡^ model (MMRT-1L, (*33*)). The Δ*C*^‡^ values determined by MMRT from these experimental measurements for cold- and meso-TIM are –4.2 ± 0.5 kJ·mol^−1^K^−1^ (at 304 K (*T*_0_)) and –1.25 ± 0.07 kJ·mol^−1^K^−1^, respectively (**Figure 1B**, **Figure S1 and S2**).

**Figure 1.**
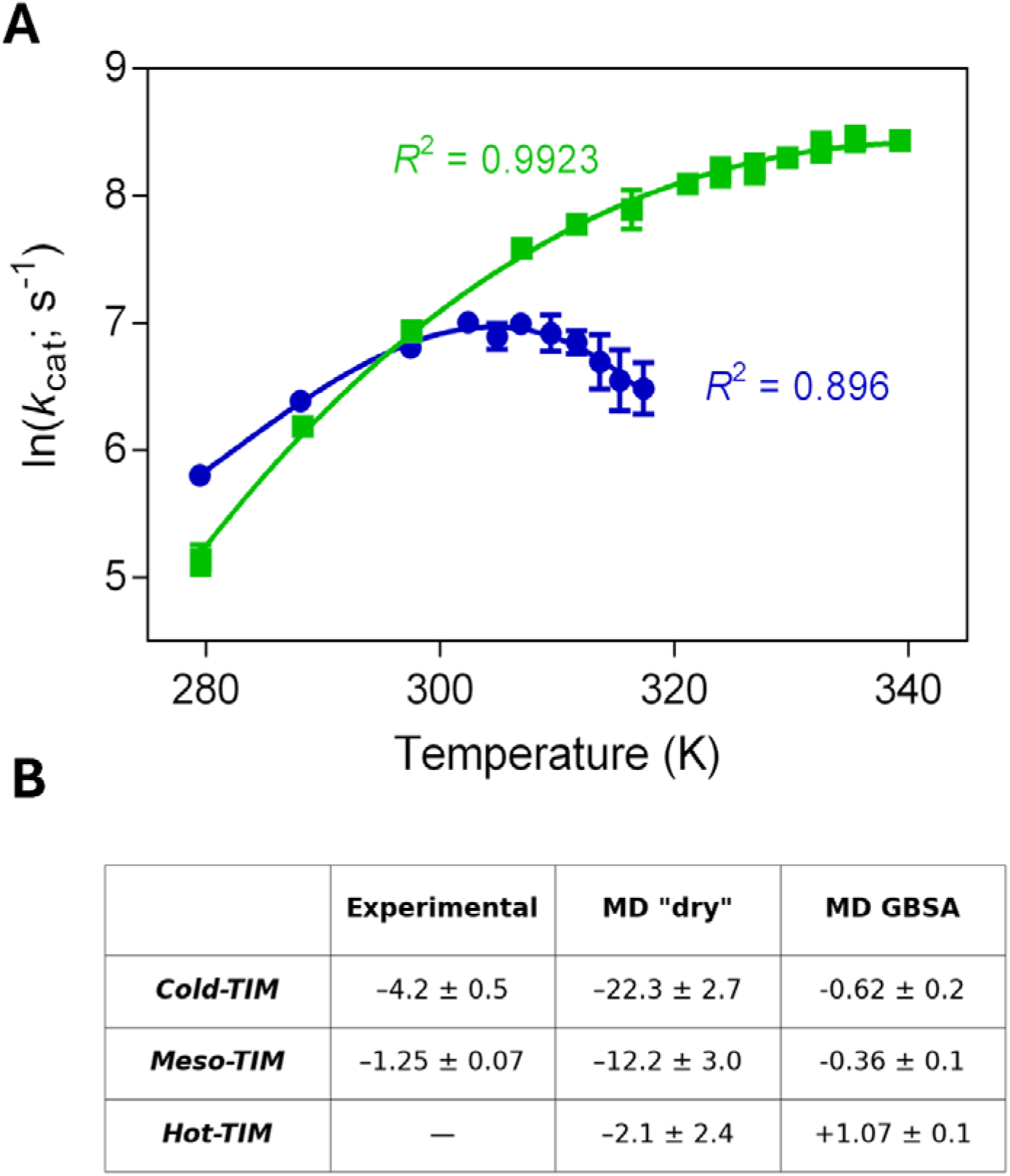
Activation heat capacity derived from (A) experimental rate-temperature measurements and (B) molecular dynamics simulations for TIM variants. **(A)** Experimentally determined temperature dependence of rates (steady state *k*_cat_) for cold-TIM (blue circles) and mesophilic meso-TIM (green squares). Data are fit with MMRT, with a temperature-dependent Δ*C*^‡^ for cold-TIM (*33*) and temperature-independent Δ*C*^‡^ for meso-TIM (see Methods). Error bars represent the standard deviation of three repeated experiments. (B) Calculated Δ*C*^‡^ values in kJ·mol^−1^K^−1^ from MD simulations of doubly occupied dimers. Computational simulation values are shown for both ‘dry’ (protein + ligand only) and GBSA (implicit solvation) calculations.

We calculated activation heat capacities (Δ*C*^‡^) for hot-, meso- and cold-TIM, at 300K based on previously validated protocols (*7*, *35*). We tested the effects of simulation conditions and models (see Supporting Information). Δ*C*^‡^ can be estimated from MD simulations, from the difference in energy variance of the RS and TS ensembles (*7*, *35*). Here, we took the enediolate (which is similar to the TS (*24*)) as a proxy for the TS (**Figure S3 and S4)**. We simulated both singly and doubly occupied enzymes, with the TS analogue present in only one active site in each. We focus here on the doubly occupied enzymes; the singly occupied simulations give generally similar results (**Figure S6 & S8**).

The simulations show a significant difference in Δ*C*^‡^ between the TIMs: cold-TIM has the the most negative Δ*C*^‡^ and meso-TIM a significantly less negative Δ*C*^‡^, in agreement with our experimental findings; hot-TIM shows only a very small Δ*C*^‡^. These findings are in agreement with predictions from MMRT (*6*), namely: the lower the temperature optimum of the enzyme, the larger the absolute value of Δ*C*^‡^.

The energy of the TS complex fluctuates less than the RS for all three enzymes, giving a negative Δ*C*^‡^ in each case. The calculated activation heat capacities are significantly different between the three TIMs and are in the order predicted by MMRT, i.e. Δ*C*^‡^ is most negative for cold-TIM (–22.1 ± 2.7 kJ·mol^−1^K^−1^ for the doubly occupied enzyme), and smallest for hot-TIM (–2.1 ± 2.7 kJ·mol^−1^K^−1^), with meso-TIM in between (–12.0 ± 2.9 kJ· mol^−1^K^−1^). The calculated values of Δ*C*^‡^ are overestimated because they are calculated for the protein+ligand only (i.e., ‘dry’; the structures are sampled from simulations in explicit solvent, but solvent is not included in calculating the energy variance) (*4*, *5*, *7*, *36*). Direct calculation of the solvent contribution is not feasible for explicit solvent simulations because of the difficulty in quantifying variance with large numbers of water molecules. We tested effects of solvation by recalculating energies employing the generalized Born surface area (GBSA) implicit solvent model (*7*). As found for other enzymes (*35*), the resulting Δ*C*^‡^ values are lower in magnitude than for the ‘dry’ (i.e. protein + ligand only) calculations, but the trends are the same, and consistent with our experimental data and the predictions of MMRT: i.e., the absolute value of Δ*C*^‡^ is smallest for hot-TIM, and largest for cold-TIM; i.e. cold-TIM has the most negative value of Δ*C*^‡^ (**Table 1**). Δ*C*^‡^ calculated with and without implicit solvent is given in **Figure 1B** and **Figure S6 and S7** for cold-TIM, meso-TIM and hot-TIM, and their respective variance values are given in **Table S3**.

### RS and TS complexes are conformationally distinct in cold-TIM and meso-TIM but similar in hot-TIM

Analysis of global conformational behaviour for each enzyme in each state (by clustering based on Cα RMSD, see Methods and **Figure 3**) showed differences in conformational behaviour between the enzymes, consistent with the flexibility analysis. Hot-TIM samples a smaller conformational space (fewer clusters, using the same cluster cut-off) than the other two enzymes: only one major cluster is observed in both the RS and TS ensembles (covering 95% of total conformations). Meso-TIM (6 clusters with ≥4% occupancy) and (particularly) cold-TIM sample a larger conformational space (9 clusters with ≥4% occupancy); they also sample different clusters in the different (RS and TS) complexes (**Table S4**). In both cold-TIM and meso-TIM, the RS and TS sample overlapping but notably different conformations.

Cold-TIM shows the largest conformational diversity, as indicated by the highest number of clusters (9). The most populated cluster represents 17% of the trajectories (substantially less than for meso-TIM, see below); this cluster is notably dominated by conformations sampled in the TS (32.6% of all TS conformations) compared to RS (<2% of RS conformations). The opposite is the case for the next two conformational clusters which represent 27.5% and 21.4% of RS conformations and almost no TS conformations. Conversely, the next cluster is only found for TS (representing 16.4% of conformations). This pattern of distinct conformational clusters for RS and TS continues for the remaining 10 clusters (covering between 2-6% of total conformations). Only 2.5% of RS conformations (combined) are present in all clusters that represent at least 1% of TS conformations. Each of the 14 clusters is thus dominated by either the RS or the TS, indicating distinct conformational preferences in each state: the TS and.RS sample different conformations in cold-TIM.

Meso-TIM has a somewhat less diverse conformational landscape than cold-TIM, but more diverse than hot-TIM, with multiple clusters (6) with significant occupation (using the same clustering criteria). The most populated conformational cluster represents 36% of the total simulations and is prominent in both the RS (33%) and TS (38%). Notably, the next highest populated cluster is dominated by RS (31% of RS, 8% of TS), whereas the following two are predominantly sampled in TS (16% and 15% of TS, ≤ 4% of RS). This indicates multiple conformations in both states, but with distinct preferences between the states. The remaining 5 clusters, not shown here, represent between 2-8% of the conformations, with some sampled by both RS and TS and others predominantly in one (**Table S4**).

In Hot-TIM, a single cluster dominates both the RS (97%) and TS (92%), suggesting that there is only one favoured conformation along the reaction pathway at this temperature **(Figure 2)**. In other words, there is no transition between a general ES ensemble and a TS-like conformation (TLC), and thus no large heat capacity changes along the reaction coordinate are expected.

**Figure 2.**
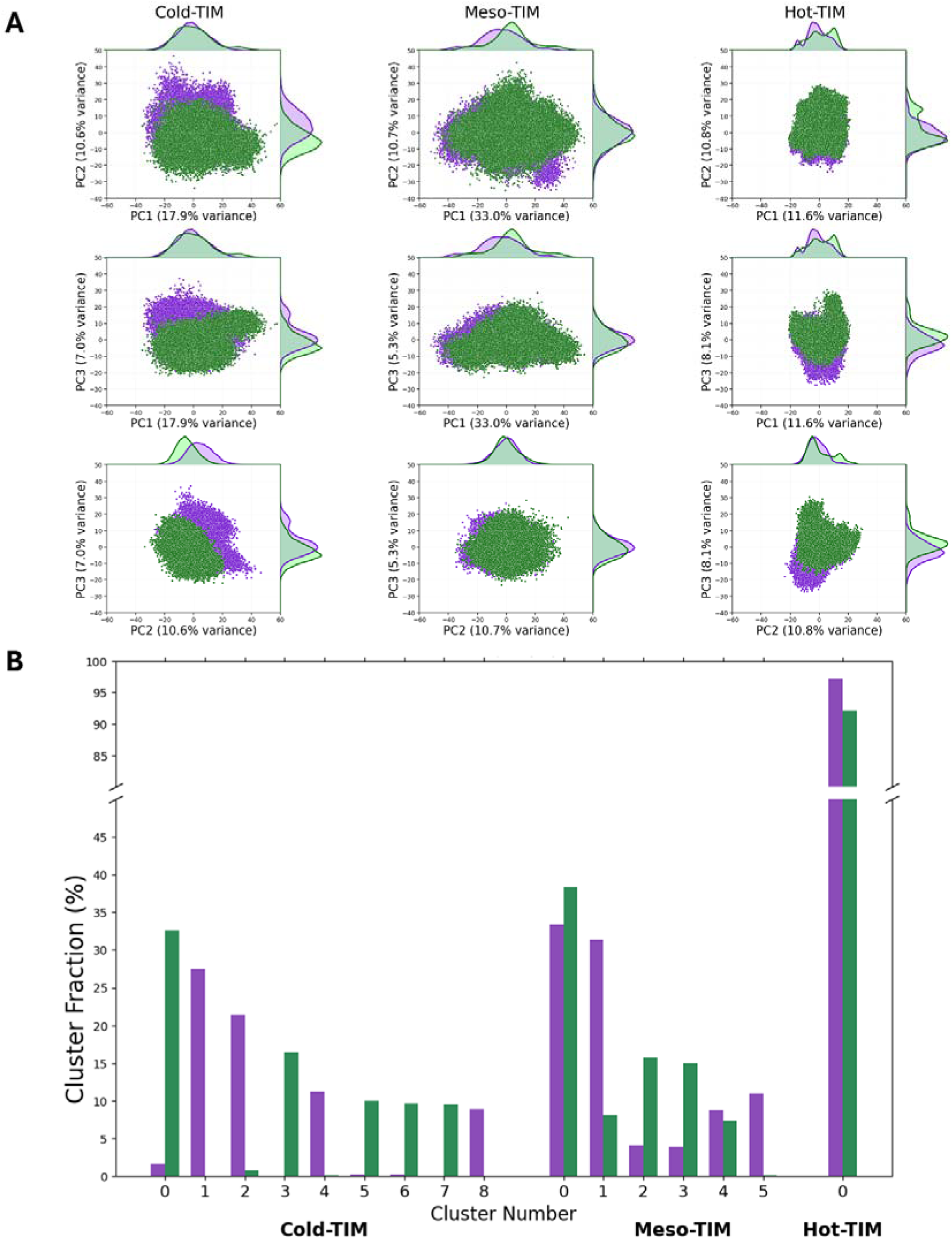
Principal component analysis and conformational clustering of TIM variants. **(A)** Principal component projections, with RS conformations in purple and TS conformations in green. Marginal density plots show the distribution along individual principal components. **(B)** “Cluster population analysis (with the same separation criterion) showing side-by-side population of each cluster for RS (purple) and TS (green) complexes.

The combination of conformational clustering and principal component analysis reveals a clear trend in how RS and TS complexes differ between the TIM homologs (**Figure 2**). Cold-TIM shows by far the largest difference in conformations between RS and TS: this shown by PCA, which shows clear separation between RS and TS ensembles, particularly along PC2. This separation indicates that the conformational transitions involve major structural rearrangements that effectively segregate the two states into distinct regions of conformational space in cold-TIM. Cold-TIM RS and TS show significant differences in conformation, indicating that a distinct, reactive conformation (TLC) is involved in the catalytic cycle.

Meso-TIM exhibits intermediate behaviour, with partial state separation evident in both analyses. PCA demonstrates that, while both RS and TS complexes explore a wide conformational landscape, PC1 shows that TS conformations are a more restricted, tighter subset of the broader RS ensemble. Like cold-TIM, meso-TIM adopts a distinct conformation in the TS, pointing to a picture in which reaction occurs in a specific TS-like conformation (TLC). This suggests that only certain RS conformations are competent for transition state formation in meso-TIM.

In contrast, the RS and TS of hot-TIM are conformationally very similar, from both types of analysis. PCA shows small absolute principal component values (indicating a small conformational space) and extensive overlap between RS and TS in principal component space, confirming that hot-TIM maintains essentially the same constrained conformational ensemble throughout the reaction at this temperature coordinate. This conformational invariance is consistent with the enzyme being pre-organized for catalysis (albeit potentially sub-optimal at this temperature) without significant structural reorganization between states.

### Change in flexibility between RS and TS is largest for cold-TIM

We analysed structural fluctuations by calculating Cα RMSF over 10 independent 500 ns simulations per state to quantify local residue-level flexibility during catalysis. Although RMSF cannot capture global conformational transitions or correlated motions (as it measures fluctuations around an average structure), it can effectively identify flexible regions within conformational basins (useful for comparing local dynamics between temperature-adapted homologs) and, particularly, changes in dynamics between different states of the same protein. RMSF analysis was performed using 10 ns windows to capture local flexibility within conformational basins while minimizing artifacts from transitions between states, and on longer timescales (450 ns) to distinguish persistent flexibility from transient fluctuations (**Figure 3, Figures S10-S12)**.

**Figure 3.**
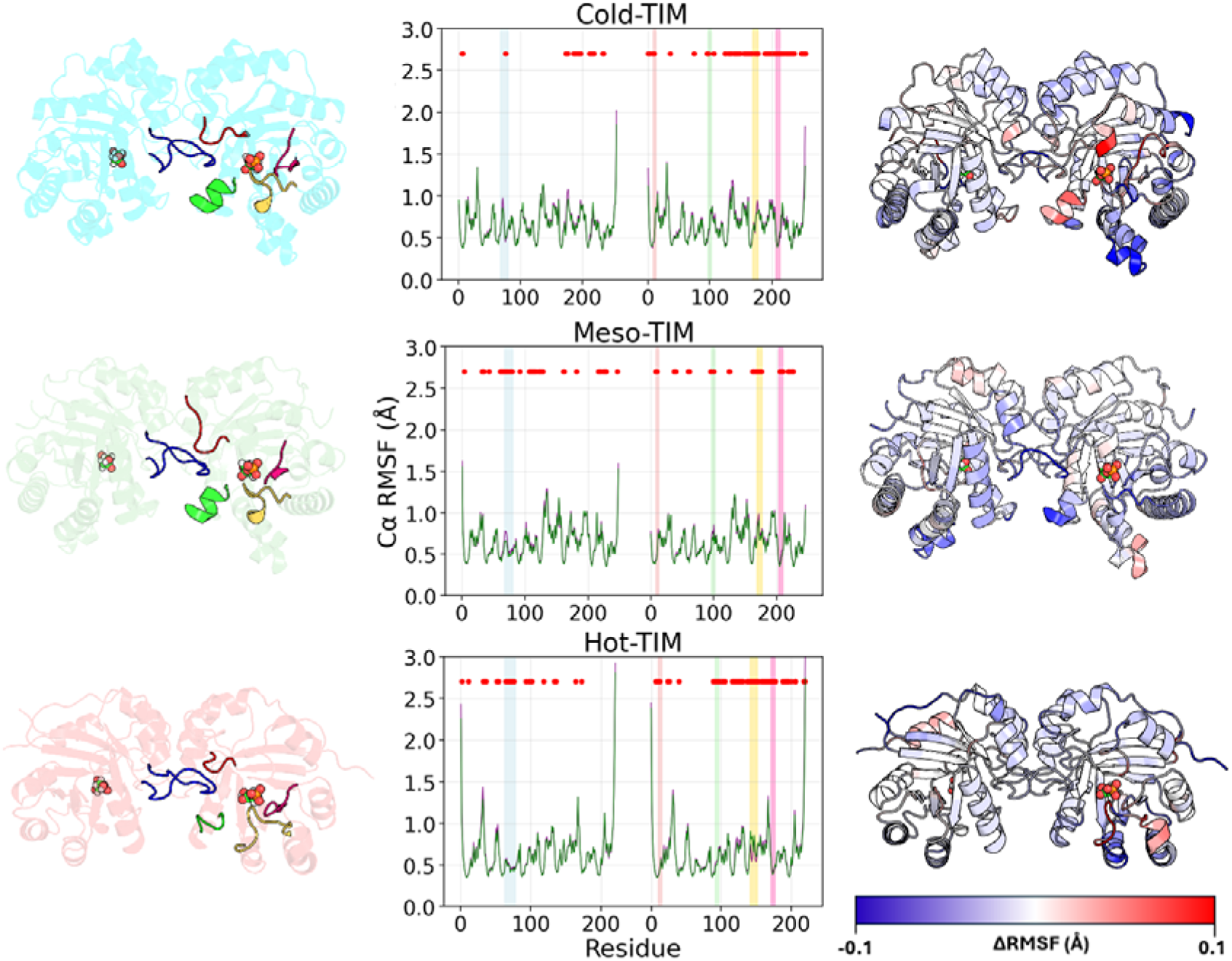
Structural fluctuations of RS and TS complexes for each TIM. Results for the doubly occupied dimers are shown here. (**left**) Structural representations showing the overall enzyme architecture with active sites highlighted (by substrate depicted in spheres). Colored motifs represent the five key dynamic regions (motifs 1-5) identified. (**middle**) Average RMSFs for RS and TS in each TIM. (Purple: RS; Green: TS). Residues for which the ΔRMSF is significant (*p* < 0.05 as determined by a two-sample t-test) are indicated by red dots. Colored bars above the plots correspond to motifs 1-5 shown in the left panel. The reacting subunit is on the right-hand side of the plot. (**right**) Cα ΔRMSF values shown on average structures. The color scale bar shows ΔRMSF units in Å, with blue indicating increased rigidity in the TS complex, and red indicating higher flexibility of the TS complex, compared to RS complex. The reacting subunit is on the right-hand side of the structure.

The magnitude of flexibility changes between states follows a clear gradient. At 450 ns timescales, cold-TIM shows mean |ΔRMSF| = 0.252 Å for residues that show significant differences in RMSF (p < 0.05, two-sample t-test, see Supporting Information). In contrast, meso-TIM shows mean |ΔRMSF| = 0.093 Å, and hot-TIM 0.087 Å (for residues with significant changes in RMSF). This is a 4.5-fold difference between cold-TIM and hot-TIM in the flexibility change between RS and TS complexes). These changes are concentrated in five conserved structural motifs that are functionally important in TIM catalysis (Figure 2): the dimer interface loop (motif 1), a conserved lysine-containing loop connecting the interface to the active site (motif 2), the HSERR helix-loop region (motif 3), loop 6 (motif 4), and loop 7 (motif 5). Despite structural conservation of these motifs across variants, they exhibit qualitatively different flexibility changes between RS and TS complexes in the different enzymes, despite all motifs being of near identical length (with the exception of motif 3 in hot-TIM).

In cold-TIM, some regions become more flexible in the TS complex while others become substantially more rigid. At short timescales (10 ns), active site motifs 2, 3, and 5 show increased flexibility in the TS. Catalytic loop 6 (motif 4) is more flexible at 10 ns, but at 450 ns timescales (capturing conformational transitions) this loop shows significant net rigidification (p < 0.05), indicating that it becomes conformationally restricted in the TS complex. Compensating for enhanced active site dynamics, surface helices (E131-G139, Q156-G163, E201-N206) and N/C-terminal regions undergo substantial rigidification in the TS. This bidirectional pattern indicates that cold-TIM employs enhanced active site flexibility to facilitate catalysis at lower thermal energy but pays an entropic cost through reduced flexibility in distal regions. Cold-TIM’s inherent flexibility means that by 300 K, active site regions (particularly loop 6) begin to exhibit excessive flexibility. Compensatory rigidification of distal surface regions maintains activity but results in a large entropic penalty, explaining why cold-TIM’s activity plateaus near 300 K. Meso-TIM’s greater inherent rigidity does not require such compensation, allowing activity to continue increasing well beyond this temperature.

Meso-TIM shows modest flexibility changes uniformly distributed across the five motifs (mean |ΔRMSF| = 0.093 Å at 450 ns). These changes are of mixed sign and smaller magnitude than cold-TIM, with no dramatic bidirectional pattern. The helices present in cold-TIM (E131-G139, Q156-G163, E201-N206) exist in meso-TIM but show no significant state-dependent changes. Moderate structural rigidification is concentrated around the active site and dimer interface, representing an intermediate adaptation strategy.

Hot-TIM exhibits minimal flexibility changes between states (mean |ΔRMSF| = 0.087 Å at 450 ns), comparable to meso-TIM in magnitude. The regulatory helices observed in cold- and meso-TIM are structurally absent in hot-TIM. An exception occurs in catalytic loop 6 (motif 4), which shows increased flexibility in the TS at both short and long timescales—opposite to the pattern in cold-TIM. This anomaly may reflect limitations in simulating the rigid thermophilic enzyme below its optimal temperature or structural accommodation issues with the transition state analogue (see Discussion).

The gradient in flexibility reorganization: cold-TIM showing extensive bidirectional changes (0.252 Å), meso-TIM and hot-TIM showing minimal changes (0.093 Å and 0.087 Å, respectively) provides a structural basis for understanding the differences in temperature adaptation across these variants. These findings are consistent with simulations of cold- and meso-TIM by Åqvist (*17*), who identified enhanced surface flexibility in psychrophilic variants. Our results extend these observations by demonstrating that flexibility changes in cold-TIM are bidirectional and state-dependent: enhanced dynamics in active regions are compensated for by rigidification in distal regions, and this pattern is fundamentally different from mesophilic and thermophilic homologues.

### Changes in correlated dynamics between TS and RS complexes are more pronounced in Cold-TIM

To analyse dynamical correlations between RS and TS complexes, dynamical cross correlation matrices (for Cα atoms) and corresponding shortest path maps (SPMs) (*37*) were calculated for each state in each enzyme. SPMs identify which residues move together in a coordinated fashion, revealing how conformational changes propagate through a protein structure. Network topology was quantified using established graph-theoretic metrics: Jaccard similarity (fraction of shared connections between states), betweenness centrality (how critical a residue or connection is for propagating motion through the structure), and normalized mutual information (similarity of modular organization).

18% of residue pairs in cold-TIM show increased correlation and anti-correlation (|Δ| > 0.1; **Figure 4**) between RS and TS, compared to 11% in meso-TIM and only 1.9% in hot-TIM. This pattern persists across all thresholds (|Δ| > 0.05, 0.2, 0.3, or 0.4) (**Table S5**): cold-TIM consistently shows the highest percentage of correlation increases across different thresholds, and hot-TIM the lowest. With a stringent threshold of |D| > 0.4, cold-TIM still shows frequent correlation increases (0.2%) within the reacting subunit and across the dimer interface, extending to the second active site (**Figure 4**). Meso-TIM still has some significant changes (0.08% of pairs) at this threshold, primarily across the dimer interface, whereas no correlation changes > 0.4 are observed in hot-TIM.

**Figure 4.**
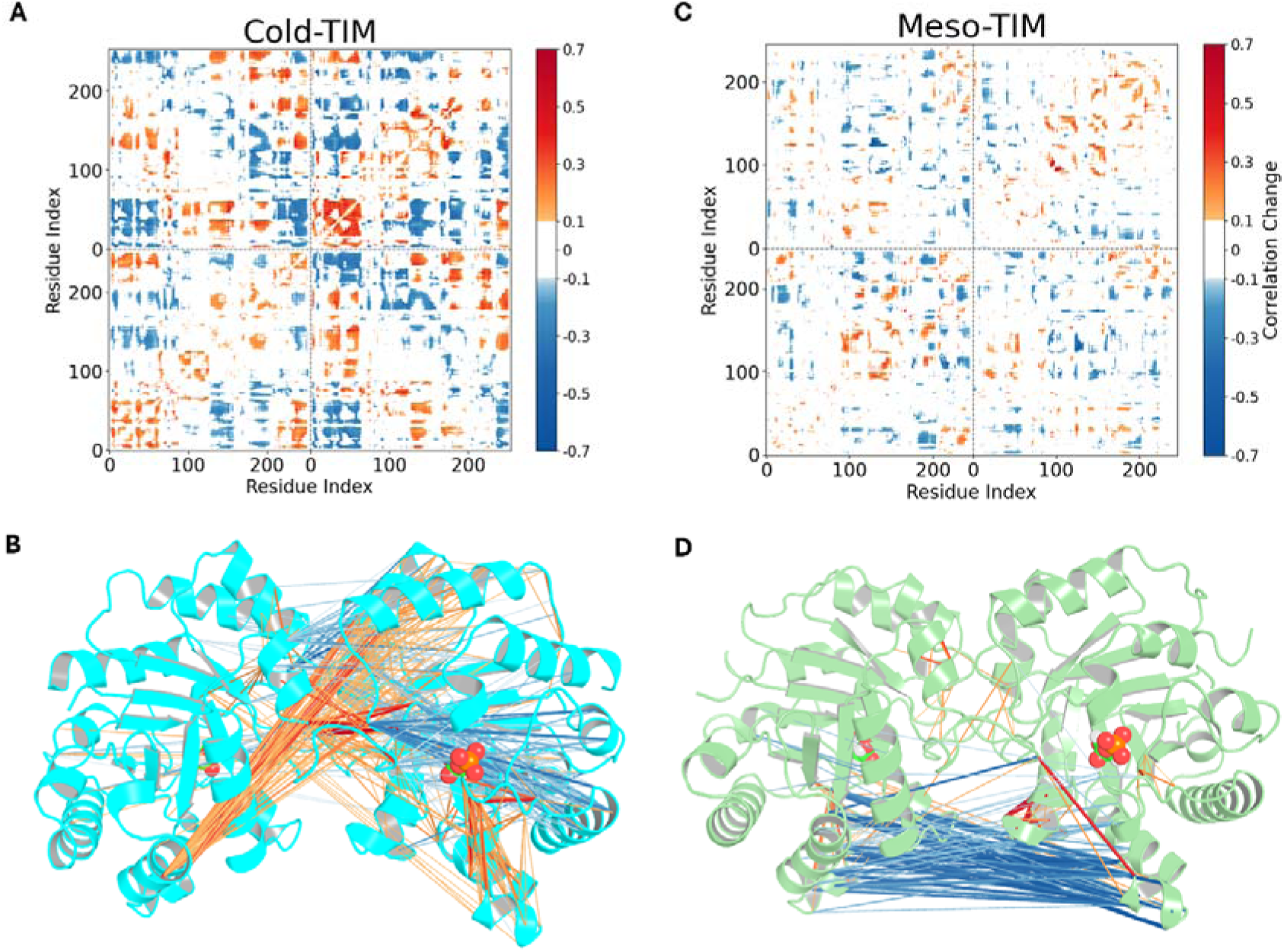
Changes in correlated dynamics between RS and TS complexes in cold-TIM and meso-TIM. Difference matrices show changes in correlation (TS-RS) for (A) cold-TIM and (C) meso-TIM. Only changes with magnitude greater than 0.1 are shown, with blue representing increased correlation and red representing increased anti-correlation. Dashed black lines indicate the monomer sequence boundary. Lower panels B and D show these correlation changes mapped onto protein structures for (B) cold-TIM and (D) meso-TIM, showing only changes above a threshold of 0.4. Red/orange lines indicate increases in correlation, while blue lines show increases in anti-correlation.

Differences observed in the correlation patterns are also apparent from the corresponding SPMs, which reveal differences in communication networks both between RS and TS complexes, and across TIM variants. SPMs can help to identify differences in structural dynamics and highlight positions that act as nodes within a dynamic network. SPM analysis reveals distinct patterns of coordinated motion (**Figure 5**). In cold-TIM, the residues that move together change almost completely between RS and TS. The RS network (25 nodes, 22 connections) and TS network (17 nodes, 16 connections) share only 18% of their connections; only 6 residues participate in coordinated motion in both states. These 6 shared residues undergo a striking transformation: their structural importance for propagating motion increases 75-fold from RS to TS (betweenness centrality: 0.007 to 0.529 for residues A41-P43). In the RS, residues A41-P43 form weak, peripheral connections; in the TS, movements throughout the enzyme propagates through these positions to other regions. This transformation provides further evidence of a TLC: occasionally in the RS, the enzyme samples conformations where these residues undergo coordinate motion like they do in the TS complex. The residues that serve as primary coordinators completely change as well: in RS, residues N67-D69 and M80 in the interface (loop 3) coordinate motion; in TS, this role shifts to residues A41 and P42 (hub position correlation *r* = –0.034, indicating no relationship between the important residues in each state). The TS coordination pattern involves a continuous path of principally hydrophobic residues that connects β2 in the reacting monomer to β2 in the non-reacting monomer through interface residues A44 and L45’ (with ‘ denoting the non-reacting monomer).

**Figure 5.**
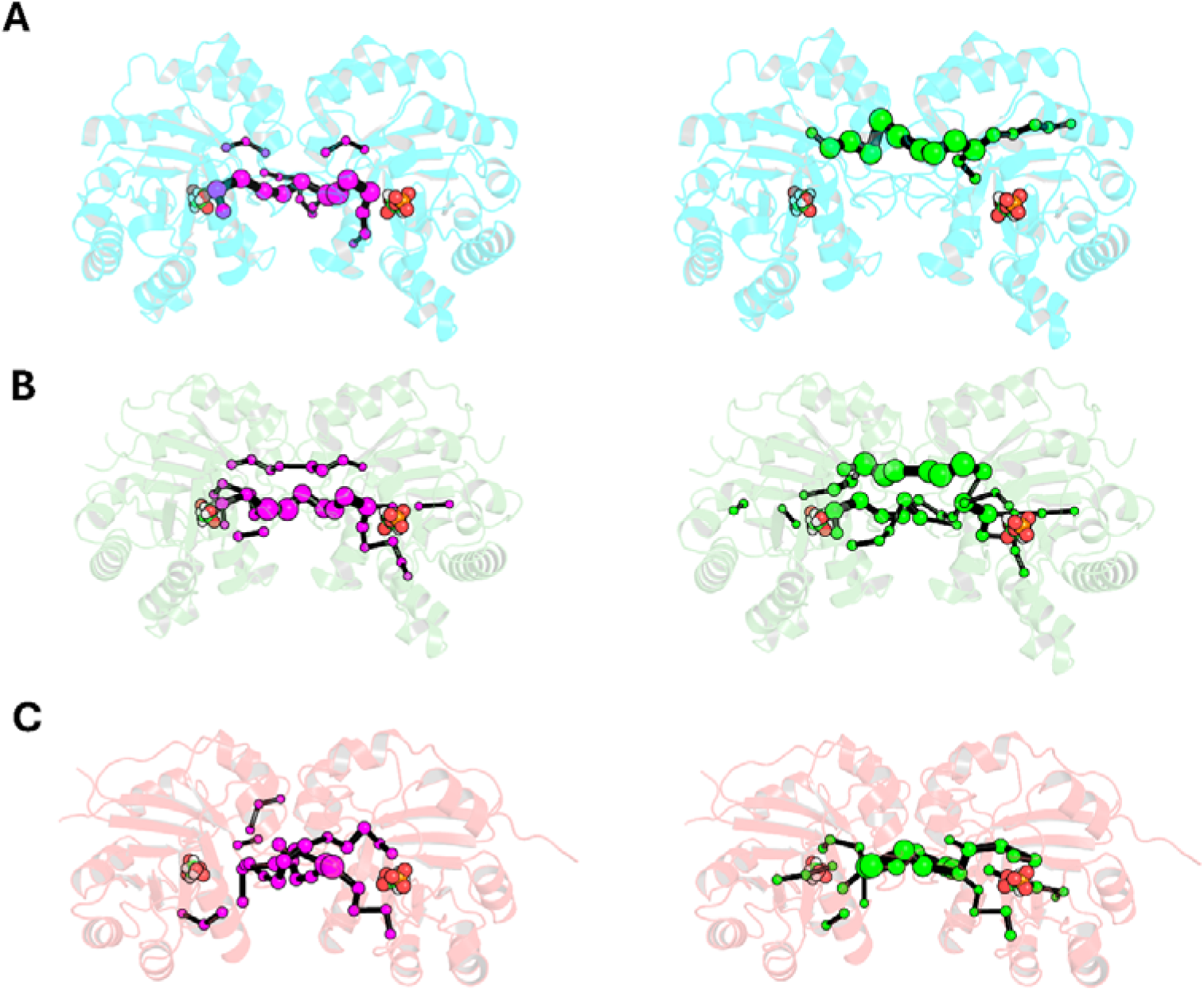
Shortest Path Maps (SPMs) of TIM variants. SPMs for (A) cold-, (B) meso-, and (C) hot-TIM variants showing reactant state (RS, left panels, magenta) and transition state (TS, right panels, green) complexes. Protein structures are shown as transparent cartoons with substrate molecules as spheres. The reacting subunit is on the right. Nodes (spheres) represent key residues in the communication network, with their size indicating relative importance, connected by edges (black lines) that represent communication pathways between residues.

In meso-TIM, many of the same residues move together in both states, but specific pathways become much more important in the TS. The RS network (32 nodes, 28 connections) and TS network (41 nodes, 36 connections) share 64% of their connections, more than three-fold greater conservation than cold-TIM. About two-thirds of the coordinated motions present in RS persist into TS. However, certain connections, particularly those involving residues C41-A44, become 6-15 times more important for coordinating motion across the enzyme (betweenness centrality: 0.014-0.030 to 0.179-0.228). Notably, the pathway containing these residues is almost identical to that in the TS network from cold-TIM, connecting β2 strands from both monomers (**Figure S13**). Rather than its coordination pattern entirely between RS and TS, meso-TIM adds new pathways (expanding from 28 to 36 connections) while strengthening specific routes. The correlation in connection importance between states is modest (*r* = 0.284), reflecting selective strengthening concentrated at the dimer interface, especially in the β2-β2**’** motif, which shows enhanced coordination. While hub positions show complete turnover (5 lost, 5 gained), their locations show moderate correlation (*r* = 0.435), indicating that new coordinators emerge in structurally related regions.

In hot-TIM, the same residues coordinate motion in the same way in both RS and TS complexes. The RS network (33 nodes, 28 connections) and TS network (34 nodes, 29 connections) differ in only 32% of specific connections. The critical metric is that the relative importance of connections for coordinating motion is nearly perfectly preserved (betweenness correlation *r* = 0.942). The residues most important for coordinating dynamics are nearly identical between states (residues H61-V62 and H73-V75 in loop 3), and their importance barely changes (maximum betweenness change: 0.014—two orders of magnitude smaller than cold-TIM). While some specific connections change, the overall pattern of motion through the protein is essentially unchanged.

Analysis of correlation loss with different cutoff thresholds showed comparable patterns in all three TIM variants (**Table S6**). At a threshold of 0.1, residue pairs losing correlation represented 1.9% (cold), 3.2% (meso), and 1.3% (hot). While the mesophilic enzyme showed slightly higher values, the overall similarity and magnitude in loss of correlation confirms that the observed increases in correlation represent genuine differences in dynamic behaviour between RS and TS, rather than redistribution of correlation patterns.

## Discussion

High-resolution kinetic measurements reveal clear non-Arrhenius behaviour for cold- and mesophilic TIMs. This curvature cannot be attributed to unfolding or a change in the rate-limiting step but instead conforms to Macromolecular Rate Theory (MMRT), which introduces, the activation heat capacity (Δ*C*^‡^) to describe how activation enthalpy and entropy vary with temperature. Both experimental and computational analyses identify Δ*C*^‡^ as a central thermodynamic parameter governing temperature adaptation: the psychrophilic enzyme exhibits the most negative value for Δ*C*^‡^, the mesophilic variant an intermediate value, and the thermophilic enzyme a near-zero value for Δ*C*^‡^, fully consistent with MMRT predictions (*4*, *6*).

Molecular-dynamics (MD) simulations conducted at 300 K, above the natural growth optimum for cold-TIM and below that for hot-TIM, show that Δ*C*^‡^ arises from differences in both conformational sampling and dynamical behaviour between the reactant-state (RS) and transition-state-like (TLC) complexes. Cluster and principal-component analyses reveal that cold-TIM explores a broad ensemble in the RS complex and samples a distinct, different conformational ensemble in the TLC, with clear separation in principal component space. The dominant TS cluster represents 32.6 % of TS conformations but fewer than 2 % of RS conformations, indicating conformational selection: the enzyme must access a specific, catalytically competent transition-state-like conformation (TLC) to react. Meso-TIM shows partial overlap between RS and TS ensembles, whereas hot-TIM remains essentially pre-organized in a reactive-like conformation at this temperature, with a single cluster accounting for 97 % of RS and 92 % of TS conformations.

Flexibility differences between RS and TS complexes form a clear gradient. All three enzymes show reduced flexibility in the TS complex compared to the RS complex, but the magnitude of this change differs substantially. Cold-TIM shows mean |ΔRMSF| = 0.252 Å for residues with significant changes (p < 0.05), compared with 0.093 Å for meso-TIM and 0.087 Å for hot-TIM. Five conserved motifs in cold-TIM display enhanced active-site flexibility in the TS complex coupled with rigidification of distal surface regions. Loop 6 becomes conformationally restricted in the TS complex at long timescales despite being flexible over short timescales. Meso-TIM exhibits moderate, mixed-sign flexibility changes, whereas hot-TIM shows minimal differences except for loop 6, which becomes more flexible in the TS complex, the opposite pattern to cold-TIM. This behaviour could reflect limitations of simulating the rigid thermophilic enzyme below its physiological temperature, imperfect accommodation of the enediolate transition-state analogue, or genuine local motions that contribute little to Δ*C*^‡^.

Dynamical networks (*38–40*) play an important role in preorganization and regulating the flexibility of enzymes for temperature adaptation (*4*, *15*). Analysis of correlated dynamics reveals extensive network reorganization in cold-TIM. Dynamical cross-correlation and shortest-path map analyses show that 18 % of residue pairs in cold-TIM exhibit increased correlation or anti-correlation between RS and TS complexes (|Δ| > 0.1), compared with 11 % in meso-TIM and only 1.9 % in hot-TIM. In cold-TIM, the coordination network is almost completely reorganised between states, with certain residues showing up to 75-fold increases in their importance for propagating motion through the structure. This provides structural evidence for the TLC concept, where RS conformations occasionally sample weak TS-like coordination patterns that become dominant at the barrier top. Meso-TIM retains 64 % of its connections but selectively strengthens pathways at the dimer interface that mirror the dominant TS network in cold-TIM. Hot-TIM preserves nearly identical coordination architecture, with changes two orders of magnitude smaller. Computational design studies demonstrate that such dynamical networks represent viable engineering targets for modulating enzyme temperature dependence through the mutation of distal residues (*41*).

The calculated activation heat capacity reflects both which conformations are sampled and how the protein fluctuates within them, encompassing both conformational sampling and internal dynamics. Cold-TIM’s transition between distinct RS and TS conformational ensembles, accompanied by extensive reorganization of flexibility and coordination patterns, generates large changes in energy fluctuations and thus large negative Δ*C*^‡^. This adaptation enables efficient catalysis at cold-TIM’s physiological temperature, where inherent flexibility allows efficient sampling of productive configurations without the extensive reorganization observed at 300 K. However, at 300 K—well above its organismal growth optimum—this same flexibility creates a substantial entropic penalty (large negative TΔS‡ (17)). The large negative ΔC‡ generates the strong curvature observed experimentally, with the entropic penalty contributing to the activity decline at elevated temperatures. Meso-TIM’s partial ensemble overlap with moderate reorganization yields intermediate Δ*C*^‡^. Hot-TIM’s minimal RS–TS differences produce near-zero Δ*C*^‡^, maintaining a relatively rigid structure with minimal conformational reorganization between states at 300 K (below its physiological temperature of ∼60°C). The extent and temperature sensitivity of conformational and dynamical reorganization thus determine each enzyme’s Δ*C*^‡^ and temperature dependence.

These results provide molecular-level validation of Macromolecular Rate Theory and reveal the physical basis of enzyme temperature adaptation. These findings extend previous work on psychrophilic flexibility (*17*, *18*, *41*) by demonstrating that cold adaptation involves state-dependent reorganization of both local flexibility and long-range coordination networks, rather than uniform increases in dynamics or simple changes in surface rigidity. The magnitude of Δ*C*^‡^ directly correlates with the extent of this reorganization: cold-TIM’s extensive conformational transitions, bidirectional flexibility changes, and near-complete rewiring of coordination networks produce large negative values of Δ*C*^‡^; meso-TIM’s moderate reorganization yields intermediate Δ*C*^‡^; and hot-TIM’s minimal reorganization results in near-zero Δ*C*^‡^at 300K. By revealing how differences in conformational sampling, flexibility patterns, and coordination networks between reaction states generate activation heat capacity changes, these results help to establish the molecular determinants of thermal adaptation in enzyme catalysis, with implications for the rational design of temperature-optimised enzymes.

## Methods

### Model construction

Crystal structures of TIM complexes with closed conformation of the loops around the active site were selected for MD simulations: these were hot-TIM from *Thermoplasma acidophilum* (PDB code: 5CSS) (*42*), meso-TIM from *Saccharomyces cerevisiae* (PDB: 1NEY) (*43*) and cold-TIM from *Vibrio marinus* (PDB: 1AW1) (*44*). The optimal temperature of these organisms are 60°C, 32°C and 15°C respectively. These structures were chosen based on their high resolution, completeness and conformation of the active site loops. Starting structures were constructed by replacing the DHAP with either the substrate or intermediate (**Figure S3**). RS and TS complexes were constructed for each TIM enzyme. Catalytic Glu165 (numbering according to meso-TIM) was treated as protonated only for the intermediate state in chain B. Any missing residues were added using MODELLER 9.23 (*45*, *46*). PropKa v3.1.0 (*47*) was used to predict protonation states at pH 7 (resulting in all residues being simulated in their standard protonation states, except as specified above). Histidine tautomers were determined based local hydrogen-bonding environment, resulting in His73 (hot-TIM), His184 (meso-TIM) and His188 (cold-TIM) being singly protonated on ND1 and all others on NE2

### MD simulations

MD simulations were performed with both singly occupied and doubly occupied dimer structures in both RS and TS complexes of three enzymes. All simulations and analysis were carried out using the AMBER package (http://ambermd.org/) utilizing the ff14SB forcefield (*48*) and following validated protocols for TIM (*30*). Ligand topologies and parameters were generated using the antechamber module from AmberTools 16. Partial atomic charges were calculated using the AM1-BCC method, and atom types were assigned according to the General AMBER Force Field (GAFF). The box was solvated with TIP4P water molecules (*49*) and the system was neutralized with explicit counterions explicit counterions (Na^+^ or Cl^−^). The solvent and ions were briefly relaxed, followed by minimization of the whole system with positional restraints (25 kcal·mol^−1^·Å^−2^) applied to Cα atoms and ligand).

The system was heated to 300 K after which the restraints on the Cα atoms were gradually reduced in 40ps. Then, after an initial 1ns equilibration in the NPT ensemble (1 atm), 500 ns production simulations were performed in the NVT ensemble. Production simulations were run with pmemd.cuda (with default direct-space non-bonded cutoffs) with the Berendsen thermostat and loose temperature coupling and pressure scaling (10 ps time constant each; see SI for tests and discussion of thermostats). Restraints were employed to keep the active site loop residues in place throughout the simulations (**Figure S4**), with equivalent restraints in both states. Ten independent simulations were performed for both states in each enzyme.

Heat capacity was calculated using techniques validated in earlier work (*7*), from structures at 10 ps intervals from 50–500 ns MD simulation windows with force-field energies recalculated after removing ions and all water molecules. Test calculations on the same structures with the GBSA implicit solvent model in AMBER to investigate the effect of implicit solvent on the energy variance.

### Analysis

MD simulations were analysed with CPPTRAJ (*50*). RMSFs were calculated for the backbone Cα atoms of each trajectory and are reported as the average of the ten independent trajectories in each case. RMSF was calculated using RMSD fitting to a running average coordinates from a time window of 10□ns. Analysis was performed using 10□ps snapshots from 50–500□ns of the simulations. Clustering on the Cα RMSD was performed using the hierarchical agglomerative algorithm with an epsilon value of 1.5. Principal component analysis (PCA) was performed on the Cα of every residue for each enzyme (RS and TSA complexes) combined.

Dynamic cross correlations and distance matrices were calculated with CPPTRAJ based on the backbone Cα position. The DCCM indicates the degree of correlated motion between pairs of atoms (here, the Cα carbon of each residue) during the simulations. Shortest pathway maps were calculated using recognised protocols (*51*). Dynamical networks were computed using cross correlations between Cα atoms in TS and RS. A distance cut-off was set to 6 Å, in accordance with previous work (*52*).

### Experimental characterisation

Expression systems for *Saccharomyces cerevisiae* and *Vibrio marinus* TIM proteins were set up with a N-terminal hexa-histidine tag in pET28b plasmid, in *E. coli* BL21 (DE3). Expression was carried out overnight in Luria-Bertani broth at 22 °C. Proteins were purified by immobilized metal ion affinity chromatography in 50 mM HEPES (pH 7.4) and 150 mM NaCl over a 20 mM.mL^−1^ imidazole gradient (20−1000 mM), followed by gel filtration chromatography [50 mM HEPES (pH 7.4), 150 mM NaCl, 10 mM MgCl_2_, 2.5 mM (NH_4_)_2_SO_4_] (*5*). Enzyme assays following the turnover of dihydroxyacetone phosphate were carried out in 50 mM HEPES (pH 7.4), 150 mM NaCl, 10 mM MgCl_2_, 2.5 mM (NH_4_)_2_SO_4_ on a stopped flow apparatus (TgK scientific, UK) with water bath heating via the t-pod attachment for accurate temperature control. Assays were coupled to the reduction of NAD^+^ via glyceraldehyde-3-phosphate dehydrogenase (GAPD; produced as described previously) from *E. coli*, for continuous measurement at 340 nm (*5*). For each enzyme, GAPD was confirmed to be non-rate limiting, substrate binding kinetics were determined at two temperatures, and thermal assays were carried out at saturating substrate concentration (or as close as where limitations existed) over 10 seconds to limit denaturation effects (if present at elevated temperatures). These details, along with precise compositions of all assays, are given in the supplementary.

Temperature versus rate profiles were fit with MMRT equations (2) and (3) (*33*) for mesophilic and psychrophilic TIM respectively, with the reference temperature (*T*_0_) set to *T*_opt_ – 4:

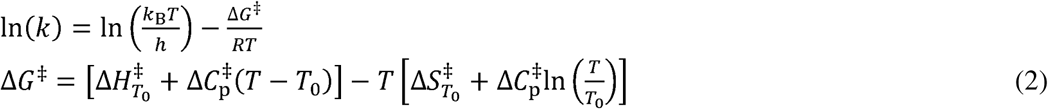

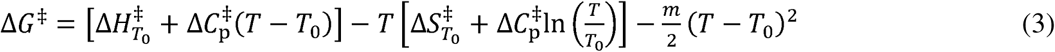

where *k* = rate coefficient; *k*_B_ = Boltzmann constant; *h* = Planck’s constant; Δ*H*^□^_*T*0_= enthalpy at *T*_0_; R = ideal gas constant; Δ*S*^□^_*T*0_ = entropy at *T*_0_. Equation (3) is an extension of (2) to incorporate linear temperature dependence of 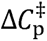, where m is the linear slope of 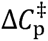. Comparison fits and statistics of equations (2) and (3) to psychrophilic data are given in the supplementary.

## Acknowledgements

This work used computational resources provided by the University of Bristol Advanced Computing Research Centre (ACRC).

## Author contributions

Conceptualization: VLA AJM MWvdK

Methodology: EJP VLA MWvdK AJM

Investigation: SS DP EJP TAW

Formal analysis: SS DP EJP

Visualization: SS DP EJP

Funding Acquistion: VLA MWvdK AJM

Writing – original draft: SS DP MWvdK EJP VLA AJM

Writing – review & editing: SS DP EJP TAW VLA MWvdK AJM

Supervision: VLA MWvdK AJM

Project administration: VLA AJM

Funding acquisition: VLA MWvdK AJM

## Data and code availability

All simulation data, including input and trajectory files, will be made openly accessible via the University of Bristol data repository, data.bris, upon manuscript acceptance.

