## Supplementary information for "Correlated dynamics, reactive conformations and non-Arrhenius behaviour in the temperature-dependence of enzyme activity: triosephosphate isomerase"

**Non-Arrhenius behaviour and dynamical differences determine thermoadaptation of triosephosphate isomerase**

### **S1. Enzyme assays**

**Optimisation of glyceraldehyde phosphate dehydrogenase (GAPDH) in assay set up**

GAPDH from *E. coli* was used as the coupling enzyme in assays to visualise assay progression through the production of NADH. This enzyme has been previously characterised under identical conditions*^1^*. Optimised cofactor concentrations were therefore taken from this characterisation, to saturate GAPDH and remain in excess over the assays. Final concentrations of 8 mM NAD^+^ and 80 mM P_i_ were used across assays.

For both TIM variants, GAPDH:TIM ratios were checked to ensure that GAPDH was in excess; that is, observed rates are equal to the rate-limiting step of TIM catalysis. Practically, this checked if doubling the TIM concentration from that used in final assays resulted in a doubling of observed rates.

**Michaelis-Menten characterisation**

Michaelis-Menten characterisations were carried out for cold-TIM at two temperatures to determine an appropriate substrate concentration for subsequent thermal characterisation (**Figure S1**). Data were fitted with an allosteric model incorporating a Hill slope (*h*) to account for cooperativity evident in the data (**equation S1**). A previously published binding constant (2.1 mM DHAP) was taken for meso‑TIM from similar characterisation conditions (25 °C, pH 7.4 (*1*)).

Equation S1	$rate=\frac{V_{max.}\left[ S \right]^{h}}{K_{\frac{1}{2}}^{h}+\left[ S \right]^{h}}$

For the fitting of cold-TIM at 30 °C, the fit is poor at low DHAP concentrations, and the poor fit results in an unrealistic Hill slope of six. A Hill slope greater than two is impossible for this dimeric enzyme (*2*). However, the data clearly shows a plateau of rates that is sufficient for determining a saturating DHAP concentration for subsequent thermal characterisation, and this anomaly was not pursued further. A DHAP concentration of 22.5 mM was utilised in the thermal characterisation of both enzymes, sufficient to largely saturate the enzyme pool for both variants.

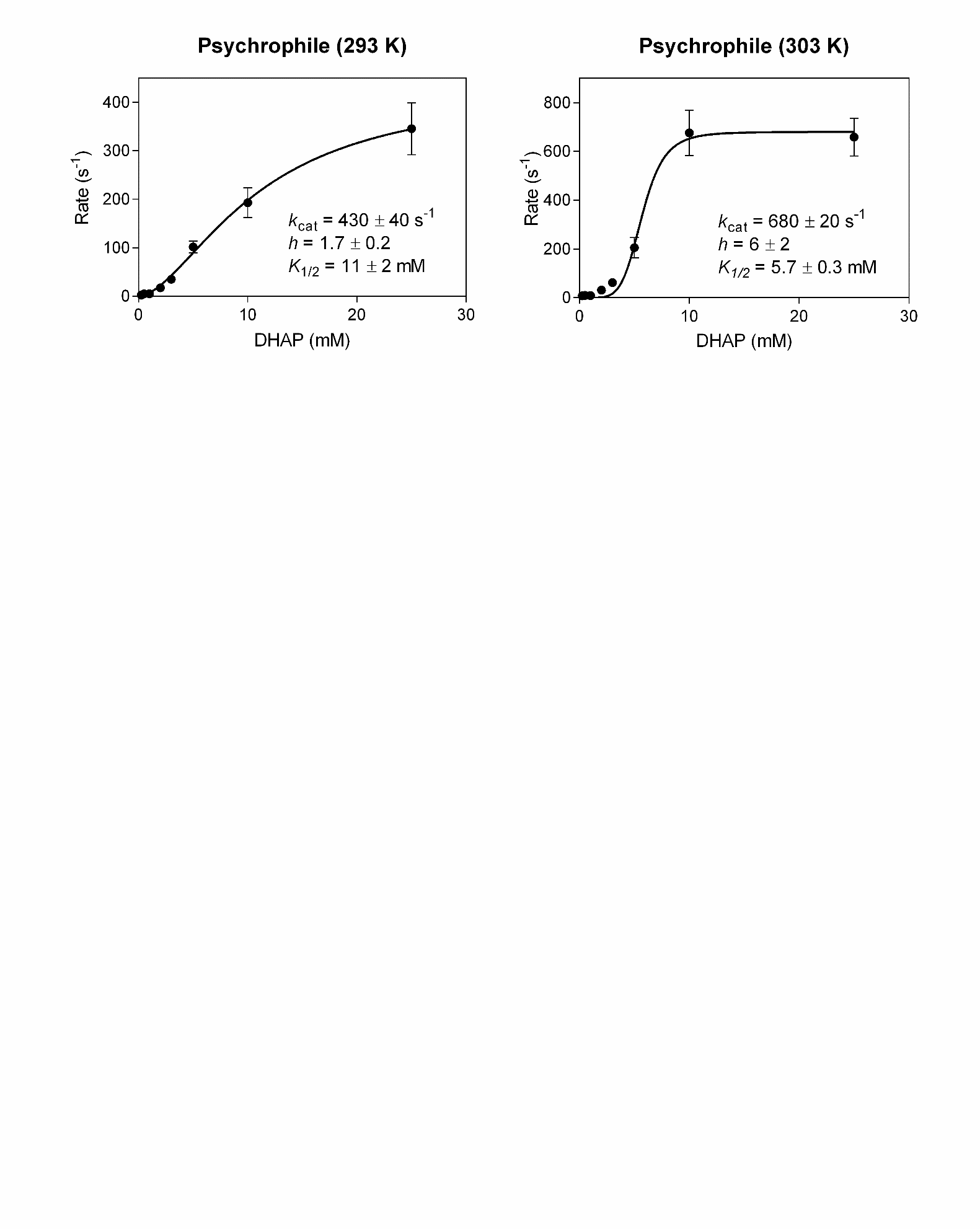

**Figure S1.** **Michaelis-Menten characterisations of cold-TIM at two temperatures**. Experimental data are fit to the Michaelis-Menten equation with the inclusion of a Hill slope (**equation S2**) to account for cooperativity.

**Assay composition details**

**Table S1.** Assay composition details for all experimental assays.

| Enzyme | Assay | Temperature | Composition |
| --- | --- | --- | --- |
| Cold-TIM | Michaelis-Menten | 293 K | Variable DHAP, 10 mM NAD^+^, 80 mM P_i_, 0.2 mg.ml^-1^ GAPDH, 0.0012 mg.ml^-1^ TPI_(psychrophilic)_ |
| Cold-TIM | Michaelis-Menten | 303 K | Variable DHAP, 10 mM NAD^+^, 80 mM P_i_, 0.2 mg.ml^-1^ GAPDH, 0.0012 mg.ml^-1^ TPI_(psychrophilic)_ |
| Cold-TIM | *T*_opt_ | - | 22.5 DHAP, 10 mM NAD^+^, 80 mM P_i_, 0.2 mg.ml^-1^ GAPDH, 0.0012 mg.ml^-1^ TPI_(psychrophilic)_ |
| Meso-TIM | *T*_opt_ | - | 22.5 DHAP, 10 mM NAD^+^, 80 mM P_i_, 0.2 mg.ml^-1^ GAPDH, 0.0005 mg.ml^-1^ TPI_(mesophilic)_ |

**Experimental Temperature dependence of** ${\boldsymbol{\Delta}\boldsymbol{C}}^{\boldsymbol{\ddagger}}$

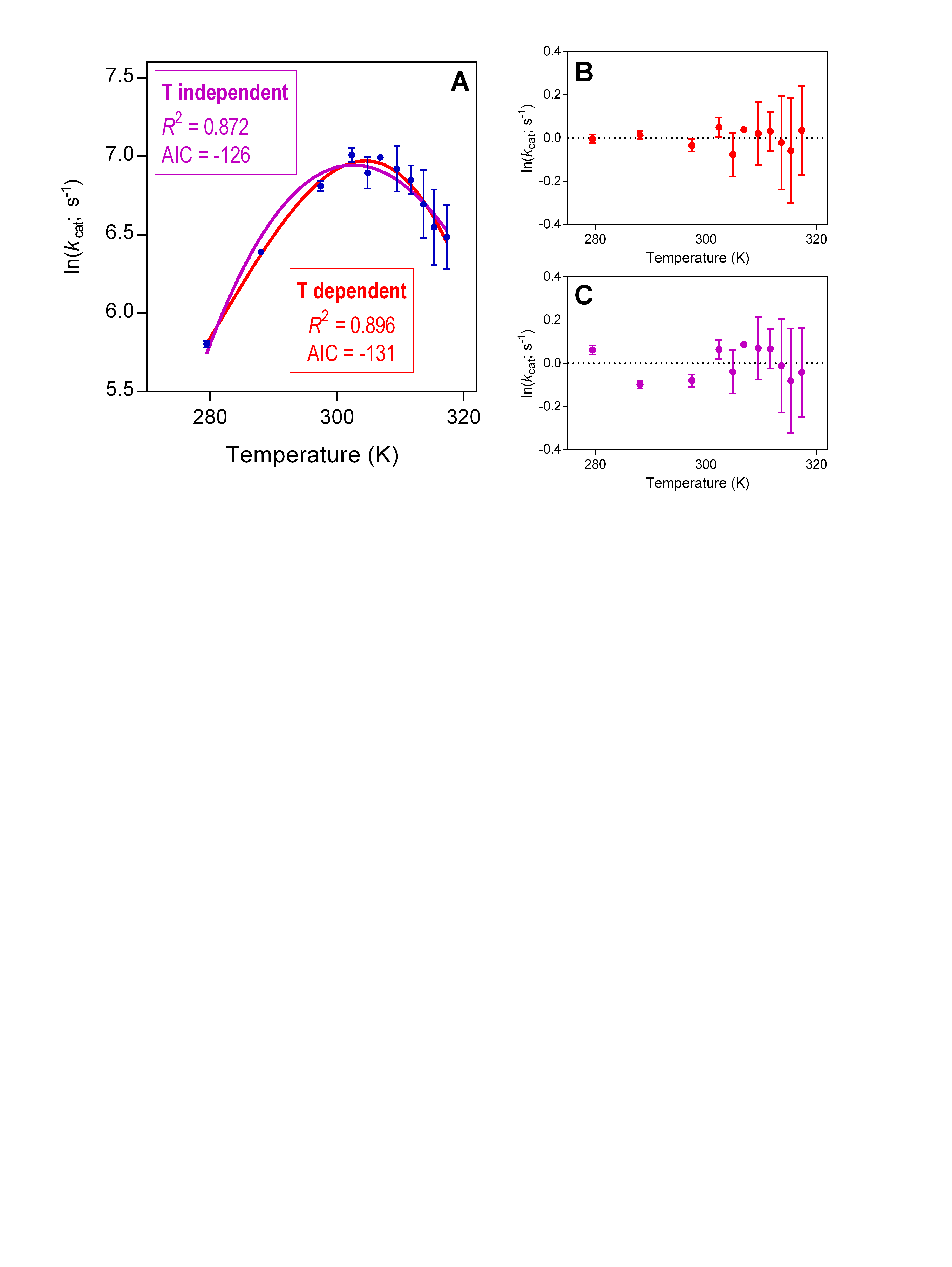

### **Figure S2.** **Comparison of cold-TIM temperature experimental kinetic data fit with (red) and without (purple) temperature dependence of** ${\boldsymbol{\Delta}\boldsymbol{C}}^{\boldsymbol{\ddagger}}$. **A**) Temperature-rate data fit with both equations and fitting statistics. **B**) and **C**) residual plots for the two fits (coloured as in (**A**)). The temperature-dependent equation (*3*) better fits the steep downturn of rates at high temperatures, and the more linear nature of the lower temperature data points. Further, the difference in Akaike information criterion **(**AIC) supports the inclusion of the extra parameter (A, the slope of the temperature dependence of ${\Delta C}^{\ddagger}$) in describing these data.

**S2.** **Reaction mechanism of TIM**

TIM catalyses isomerization of D-glyceraldehyde 3-phosphate (G3P) to dihydroxyacetone phosphate (DHAP), through a pair of enediolate reaction intermediates (*4*–*6*). Isomerization proceeds with proton transfer between carbons 1 and 2, mediated by the carboxylate side chain of Glu165 and between oxygen 1 and 2, mediated by the imidazole side chain of His95.

The crystal structures of and hot-TIM, meso-TIM and cold-TIM used for simulations (*Thermoplasma acidophilum* (PDB code: 5CSS, 440 residues) (*7*); meso-TIM from *Saccharomyces cerevisiae* (PDB: 1NEY, 494 residues) (*8*),and cold-TIM from *Vibrio marinus* (PDB: 1AW1, 509 residues) (*9*), respectively) contain bound PGA (2-phosphoglycolate), DHAP (dihydroxyacetonephosphate), and G3P (glycerol-3-phosphate), respectively. In both monomers, loop 6 is observed in a closed state in these crystal structures. For cold-TIM and hot-TIM, Pymol (*10*) was employed to place the substrate in the active site, while DHAP was manually positioned in the active site using the PGA and G3P conformations from the crystal structures as templates. In the simulations here, the DHAP-complex built from the crystal structures serves as the Reactant State (RS) complex, and the enediolate intermediate is taken as a proxy for the Transition State (TS). The corresponding ligand structures are illustrated in **Figure S3**.

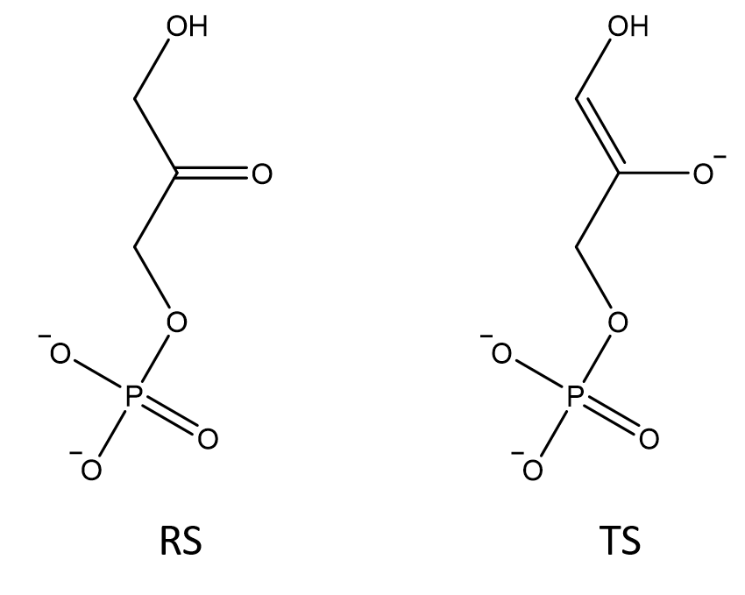

**Figure S3.** **Structure of RS and TS ligands used in simulations**. To create the TS, DHAP's conformation was adjusted using Pymol, based on the DHAP conformation. The protonation states of side chains, and histidine tautomers, were assigned based on standard pKa values in aqueous solution and visual inspection and confirmed using PROPKA 3.1(11).

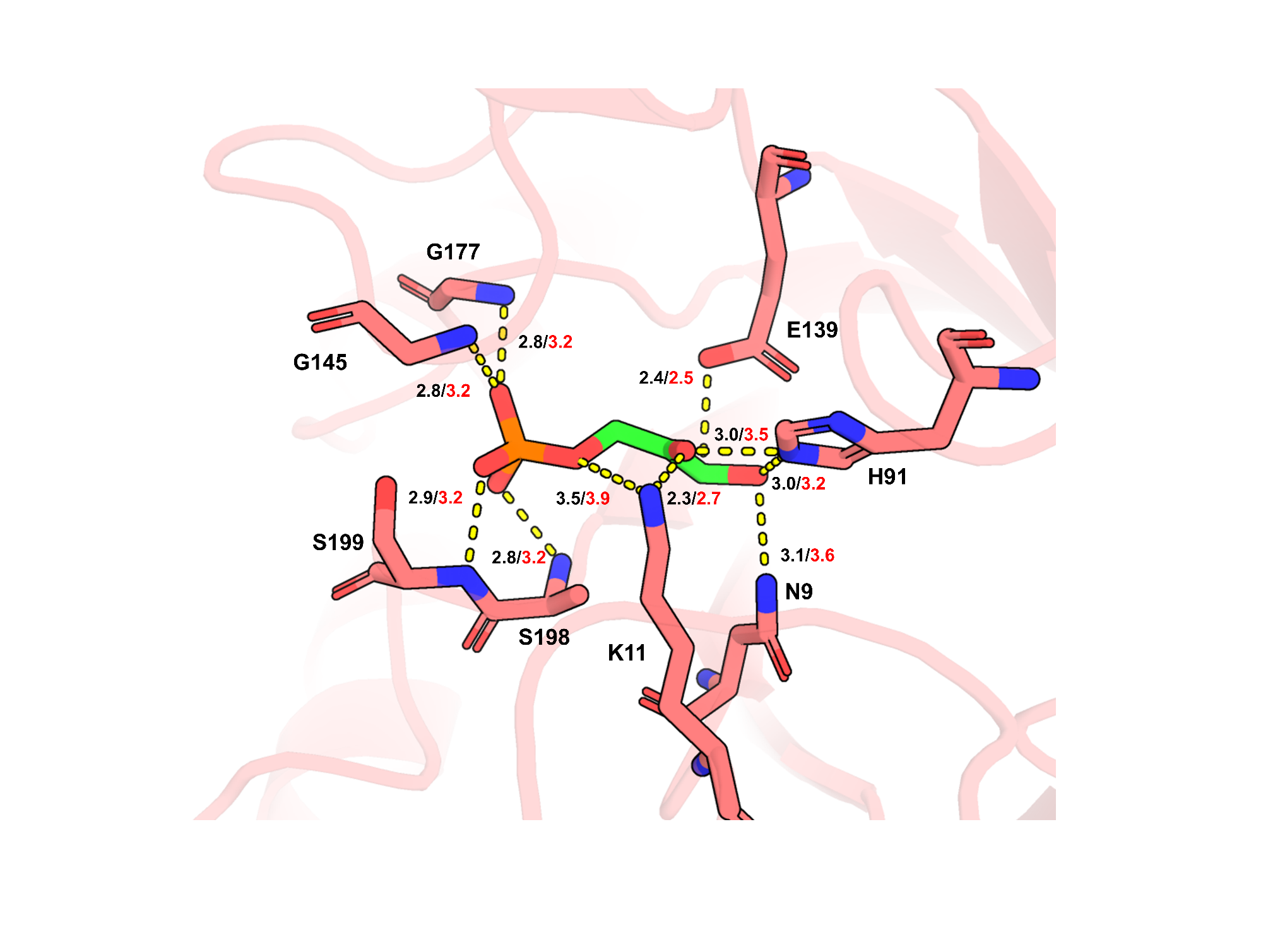

**Figure S4.** **Restraints applied in MD simulations to maintain catalytically competent conformations and prevent substrate dissociation.** Distance restraints applied to the active site of hot-TIM are shown. Black numbers indicate distances (in Å) measured in the starting crystal structure (PDB: 5CSS)(7). Red numbers show the threshold distance at which the one-sided harmonic restraint becomes active. These restraints maintain loop 6, 7, and 8 in closed, reactive conformations as observed in crystal structures of enzyme-substrate complexes. Identical restraints were applied to all three TIM variants and both ligand states to avoid biasing ${\Delta C}^{\ddagger}$calculations. Similar restraint strategies have been successfully employed in previous MD simulations of TIM(12).

TIM activity is highly sensitive to the conformation of active site loops 6, 7, and 8, which undergo substantial conformational changes during the catalytic cycle (*12*, *13*). Loop 6, in particular, transitions between open and closed conformations, with the closed state being essential for catalysis by sequestering the substrate from solvent and positioning catalytic residues (*12*). Crystal structures of enzyme-substrate complexes capture loop 6 in this closed, catalytically competent conformation. To maintain loop structures, close to those observed experimentally throughout the MD simulations, and to prevent substrate dissociation, distance restraints were applied as shown in **Figure S4** and detailed in **Table S2**. Equivalent restraints were used in each state (RS and TS) and for all three enzymes to avoid biasing the ${\Delta C}^{\ddagger}$calculations - any differences in calculated ${\Delta C}^{\ddagger}$therefore arise from intrinsic differences in conformational dynamics, not from differences in applied constraints.

Previous simulations of mesophilic TIM showed that some weak restraints are required to keep the substrate in the TIM active site during the simulations (*12*). The distances shown in **Figure S4** are the starting distances between these atom pairs in the crystal structure of hot-TIM (PDB ID: 5CSS). Starting structure for simulation (DHAP bound) with distances on which restraints are placed to keep loops 6, 7 and 8 in a reactive conformation are indicated with dashed lines. Distances in the crystal structure are shown in black, and distance where the one-sided harmonic restraint comes into effect in red (See **Table S2**). The same restraints were added to all variants and ligands to avoid biasing of the ${\Delta C}^{\ddagger}$calculations.

**Tables S2a–c. Distance restraints applied to maintain substrate positioning in TIM active sites.**

Flat-bottom harmonic distance restraints were applied to the RS and TS complexes in all three TIM enzymes to maintain active site architecture throughout the simulations. Atom nomenclature follows PDB ID: 5CSS, 1NEY and 1AW1 (residue numbers 442, 496 and 511 refer to the ligand). Parameters: r1–r4 define the flat-bottom potential boundaries (Å), with r2/r3 representing the flat-well region where no restraint force is applied; k2 and k3 are the force constants (kcal mol⁻¹ Å⁻²) for the harmonic penalties applied when distances fall outside the flat-well region. The crystal structure column reports the corresponding distances in the initial crystal structure (PDB ID: 5CSS). See **Figure S4** for visualization of the restrained atom pairs. These restraints were applied to both active sites in each system.

**Table S2a. Hot-TIM distance restraints.**

| **Atom Pair** | **r1** | **r2** | **r3** | **r4** | **k2** | **k3** | **Starting structure** |
| --- | --- | --- | --- | --- | --- | --- | --- |
| G145_N to 442_O4P | 0 | 3.2 | 3.2 | 10 | 0 | 25 | 2.8 |
| G177_N to 442_O4P | 0 | 3.2 | 3.2 | 10 | 0 | 25 | 2.8 |
| N9_ND2 to 442_O1 | 0 | 3.2 | 3.2 | 10 | 0 | 25 | 3.1 |
| K11_NZ to 442_O2 | 0 | 2.7 | 2.7 | 10 | 0 | 25 | 2.3 |
| K11_NZ to 442_O1P | 0 | 3.6 | 3.6 | 10 | 0 | 25 | 3.5 |
| A198_N to 442_O2P | 0 | 3.2 | 3.2 | 10 | 0 | 25 | 2.8 |
| S199_N to 442_O3P | 0 | 3.2 | 3.2 | 10 | 0 | 25 | 2.9 |
| H91_NE2 to 442_O2 | 0 | 3.2 | 3.2 | 10 | 0 | 25 | 3.0 |
| H91_NE2 to 442_O1 | 0 | 3.5 | 3.5 | 10 | 0 | 25 | 3.0 |
| E139_OE2 to 442_H2* | 0 | 2.5 | 2.5 | 10 | 0 | 25 | 2.4 |

**Table S2b. Meso-TIM distance restraints.**

| **Atom Pair (Meso-TIM)** | **r1** | **r2** | **r3** | **r4** | **k2** | **k3** | **Starting structure** |
| --- | --- | --- | --- | --- | --- | --- | --- |
| G170_N to 496_O4P | 0 | 3.2 | 3.2 | 10 | 0 | 25 | 2.9 |
| G210_N to 496_O4P | 0 | 3.2 | 3.2 | 10 | 0 | 25 | 2.7 |
| N9_ND2 to 496_O1 | 0 | 3.2 | 3.2 | 10 | 0 | 25 | 3.2 |
| K11_NZ to 496_O2 | 0 | 2.7 | 2.7 | 10 | 0 | 25 | 3.0 |
| K11_NZ to 496_O1P | 0 | 3.6 | 3.6 | 10 | 0 | 25 | 3.2 |
| A231_N to 496_O2P | 0 | 3.2 | 3.2 | 10 | 0 | 25 | 2.7 |
| S232_N to 496_O3P | 0 | 3.2 | 3.2 | 10 | 0 | 25 | 2.9 |
| H94_NE2 to 496_O2 | 0 | 3.2 | 3.2 | 10 | 0 | 25 | 2.6 |
| H94_NE2 to 496_O1 | 0 | 3.5 | 3.5 | 10 | 0 | 25 | 3.1 |
| E164_OE2 to 442_H2* | 0 | 2.5 | 2.5 | 10 | 0 | 25 | 2.9 |

**Table S2c. Cold-TIM distance restraints.**

| **Atom Pair (Cold-TIM)** | **r1** | **r2** | **r3** | **r4** | **k2** | **k3** | **Starting structure** |
| --- | --- | --- | --- | --- | --- | --- | --- |
| G174_N to 511_O4P | 0 | 3.2 | 3.2 | 10 | 0 | 25 | 2.6 |
| G212_N to 511_O4P | 0 | 3.2 | 3.2 | 10 | 0 | 25 | 5.6 |
| N8_ND2 to 511_O1 | 0 | 3.2 | 3.2 | 10 | 0 | 25 | 3.4 |
| K10_NZ to 511_O2 | 0 | 2.7 | 2.7 | 10 | 0 | 25 | 3.6 |
| K10_NZ to 511_O1P | 0 | 3.6 | 3.6 | 10 | 0 | 25 | 3.2 |
| G234_N to 511_O2P | 0 | 3.2 | 3.2 | 10 | 0 | 25 | 3.1 |
| G235_N to 511_O3P | 0 | 3.2 | 3.2 | 10 | 0 | 25 | 3.6 |
| H96_NE2 to 511_O2 | 0 | 3.2 | 3.2 | 10 | 0 | 25 | 2.2 |
| H96_NE2 to 511_O1 | 0 | 3.5 | 3.5 | 10 | 0 | 25 | 2.6 |
| E168_OE2 to 511_H2* | 0 | 2.5 | 2.5 | 10 | 0 | 25 | 3.0 |

*For ligand-H2: The restraint distance is measured to the hydrogen atom H2 on C1, which is added during the tleap processing step. The crystal structure distance shown is measured after adding this hydrogen to the crystal structure coordinates, representing the catalytically relevant OH distance for proton transfer.

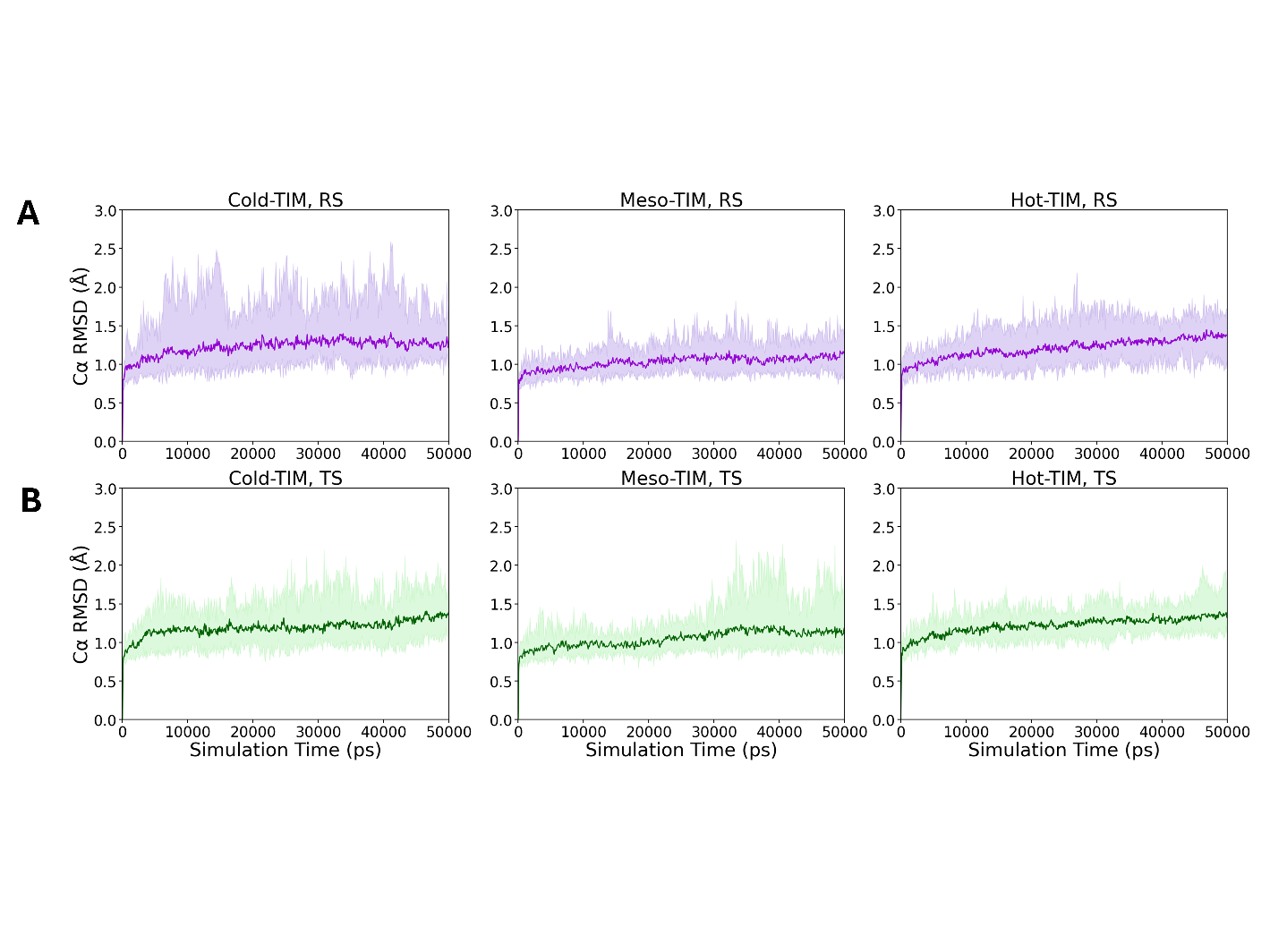

**Figure S5.** **Cα RMSD for simulations of different TIMs.** **A**) RMSD for all ten RS trajectories (purple), with the average for all 10 runs indicated by the dark purple lines. **B**) RMSD for all ten TS trajectories (green), with the average for all 10 runs indicated by the dark green lines.

**S3. Heat capacity calculations**

Normalization to a moving average was used to compute energy distributions (50 ns window), as in previous work. Heat capacities were estimated in the same way (*14*), using the moving average of the energy variance (equation 1), wher$e {\Delta C}^{\ddagger}$is the difference in heat capacity between two states, A and B with the variances $\left\langle{\delta H}_{A}^{2} \right\rangle$ and $\left\langle{\delta H}_{B}^{2} \right\rangle$. Error-bars were calculated using leave-one-out cross-validation.

${\Delta C}^{\ddagger}=\frac{\left\langle{\delta H}_{A}^{2} \right\rangle-\left\langle{\delta H}_{B}^{2} \right\rangle}{k_{B}T^{2}}$	(1)

The potential energy is calculated for the enzyme–ligand system (all atoms) as an approximation for the system enthalpy. Ten replicate 500 ns MD simulations were conducted for, both the RS and TS, a total of 5 μs of explicit solvent MD simulation for each complex, for each TIM, using protocols that we have previously tested (*12*, *14*–*16*).

**S4. Singly and doubly occupied TIM**

We tested different models by investigating two distinct types of complexes: in one, a ligand was present solely in one subunit of the dimer, while the other subunit was unoccupied (referred to here as the singly occupied dimer). The second type of complex contained a ligand in both subunits of the dimer (referred to as the doubly occupied dimer). In simulations of the doubly occupied dimer, in the RS, both active sites contain substrate; in the TS complex, one active site contains substrate and the other contains the TSA: i.e. a reacting subunit and a non-reacting subunit. This comparison tested the impact of ligand occupancy on the results.

For each simulation, the energy variance was calculated using a moving average along the 500 ns trajectory to account for transitions between conformational substates. The window size for the moving average was adjusted between 10 and 100 ns; the ${\Delta C}^{\ddagger}$ values converge around 80 ns in both doubly occupied (**Figure** S**6 and S7)** and singly occupied **(Figure** S**8**) dimers. The potential energy distribution across all three enzymes is narrower for the TS complex than for the corresponding RS complex (though not significantly so for hot-TIM), leading to a negative activation heat capacity ${\Delta C}^{\ddagger}$ (**Figure S9**).

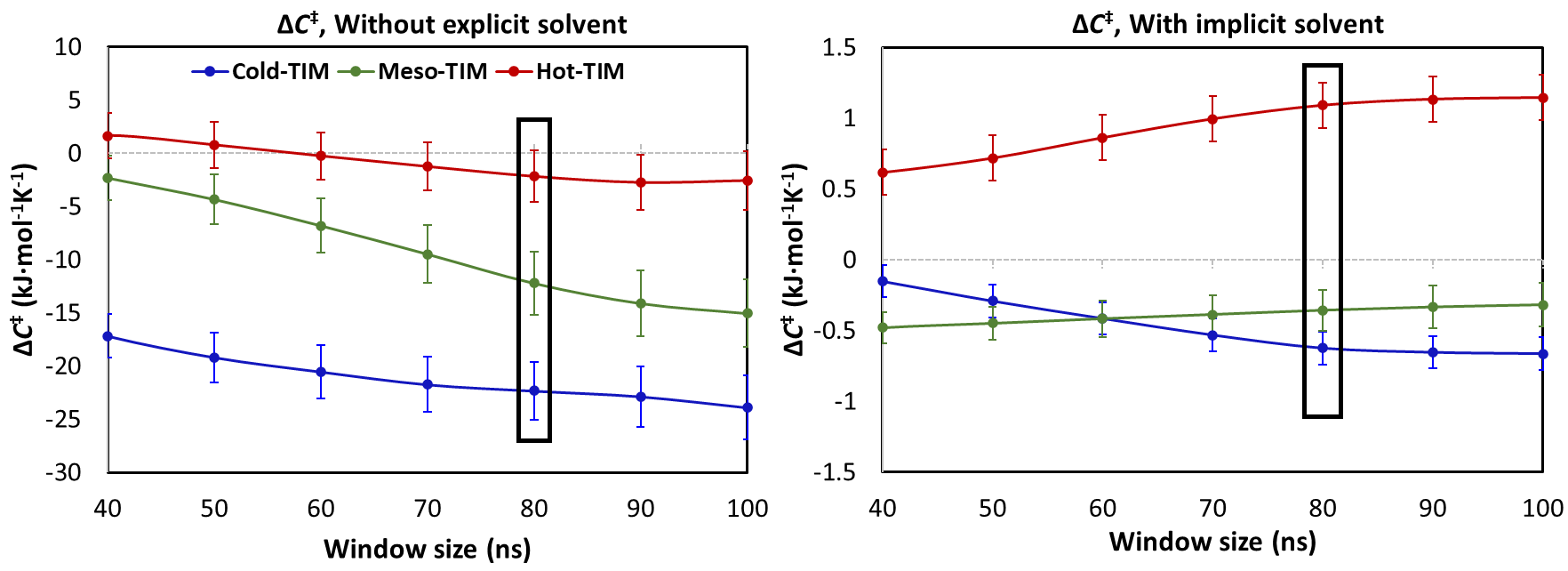

**Figure S6.** ${\Delta C}^{\ddagger}$of cold-TIM, meso-TIM and hot-TIM, from simulations of the doubly occupied dimers calculated without explicit solvent (left) and with implicit solvent (GBSA) (right). The box shows the ${\Delta C}^{\ddagger}$value at 80ns window size.

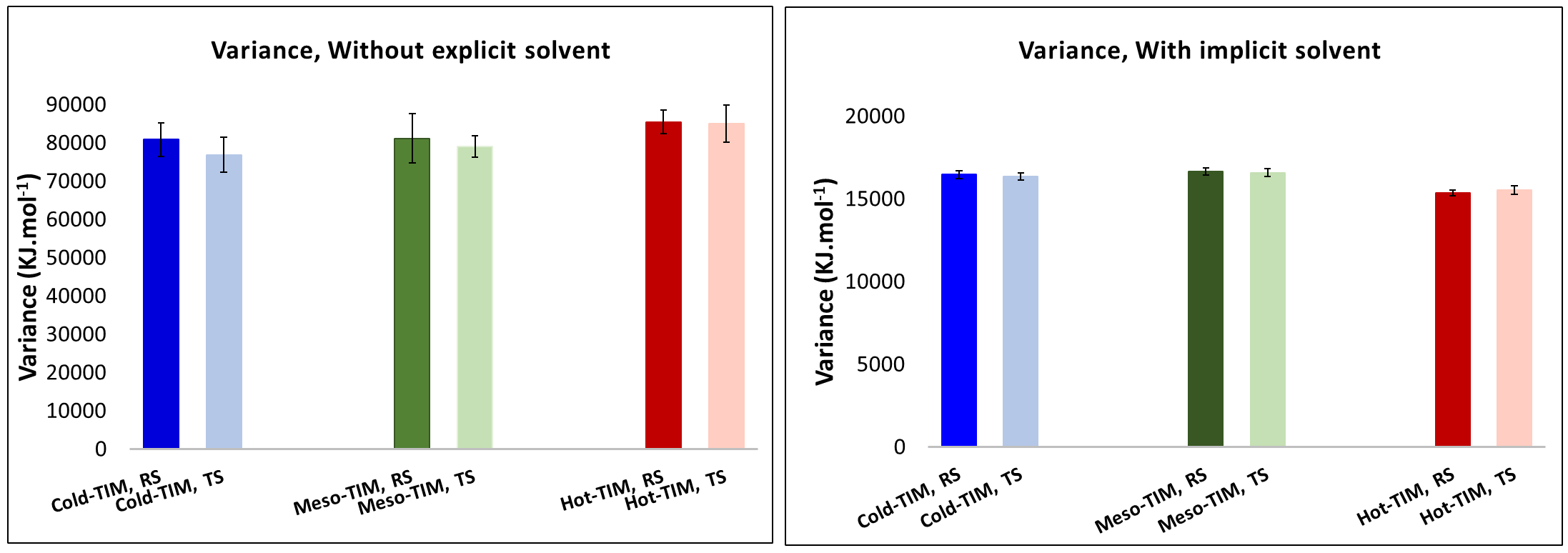

**Figure S7.** Energy variance (kJ·mol^−1^) of cold-TIM, meso-TIM and hot-TIM in RS and TS (doubly occupied dimer), without explicit solvent (left) and with GBSA implicit solvent at window size of 80 ns (right).

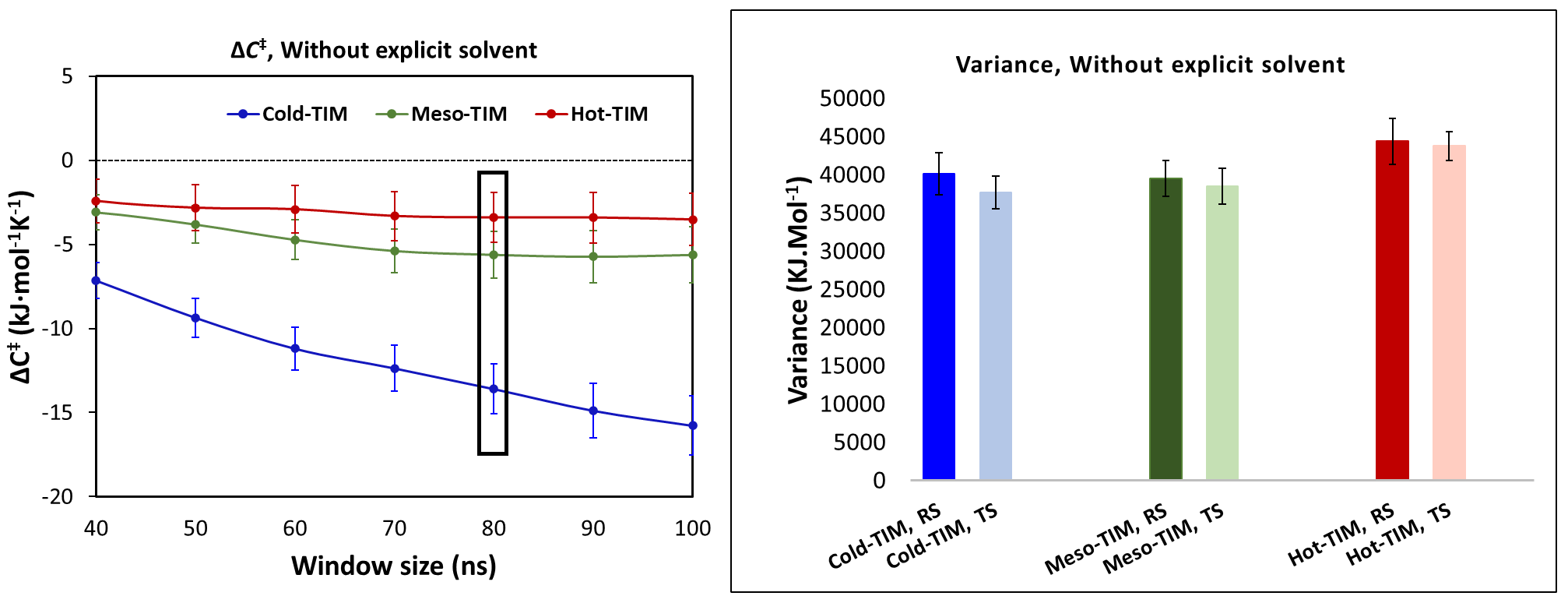

**Figure S8.** Simulations of singly occupied dimers: ${\Delta C}^{\ddagger}$ of cold-TIM, meso-TIM and hot-TIM as a function of window size (left); energy variance (kJ·mol^−1^) in RS and TS complexes in each TIM at a window size of 80ns (right) of only the reactive monomer in the singly occupied dimer.

**Table S3.** **Energy variance (kJ·mol⁻¹) in different TIM within singly and doubly occupied dimers.** The table compares energy variances across various organisms in two distinct complexes: the singly occupied dimer and the doubly occupied dimer along with their standard deviation and difference between variance values between RS and TS within each organism. For the doubly occupied dimer, calculations were conducted under two conditions: one without explicit solvent (dry state) considering only enzyme and ligand, and the other with an implicit solvent.

|  | Cold-TIM, RS | Cold-TIM, TS | Meso-TIM, RS | Meso-TIM, TS | Hot-TIM, RS | Hot-TIM, TS |
| --- | --- | --- | --- | --- | --- | --- |
| Doubly occupied dimer, without explicit solvent | | | | | | |
| Variance | 80832 | 76890 | 81178 | 79025 | 85442 | 85062 |
| Std. Dev. | 4456 | 4634 | 6432 | 2751 | 3070 | 4826 |
| **Difference** | **3942** |  | **2153** |  | **380** |  |
| Doubly occupied dimer, implicit | | | | | | |
| Variance | 16462 | 16352 | 16646 | 16582 | 15349 | 15542 |
| Error | 234 | 235 | 230 | 236 | 180 | 259 |
| **Difference** | **110** |  | **64** |  | **-192** |  |
| Reactive monomer of singly occupied dimer, without explicit solvent | | | | | | |
| Variance | 40116 | 37717 | 39507 | 38515 | 44369 | 43768 |
| Error | 2779 | 2144 | 2366 | 2346 | 2987 | 1880 |
| **Difference** | **2399** |  | **992** |  | **601** |  |

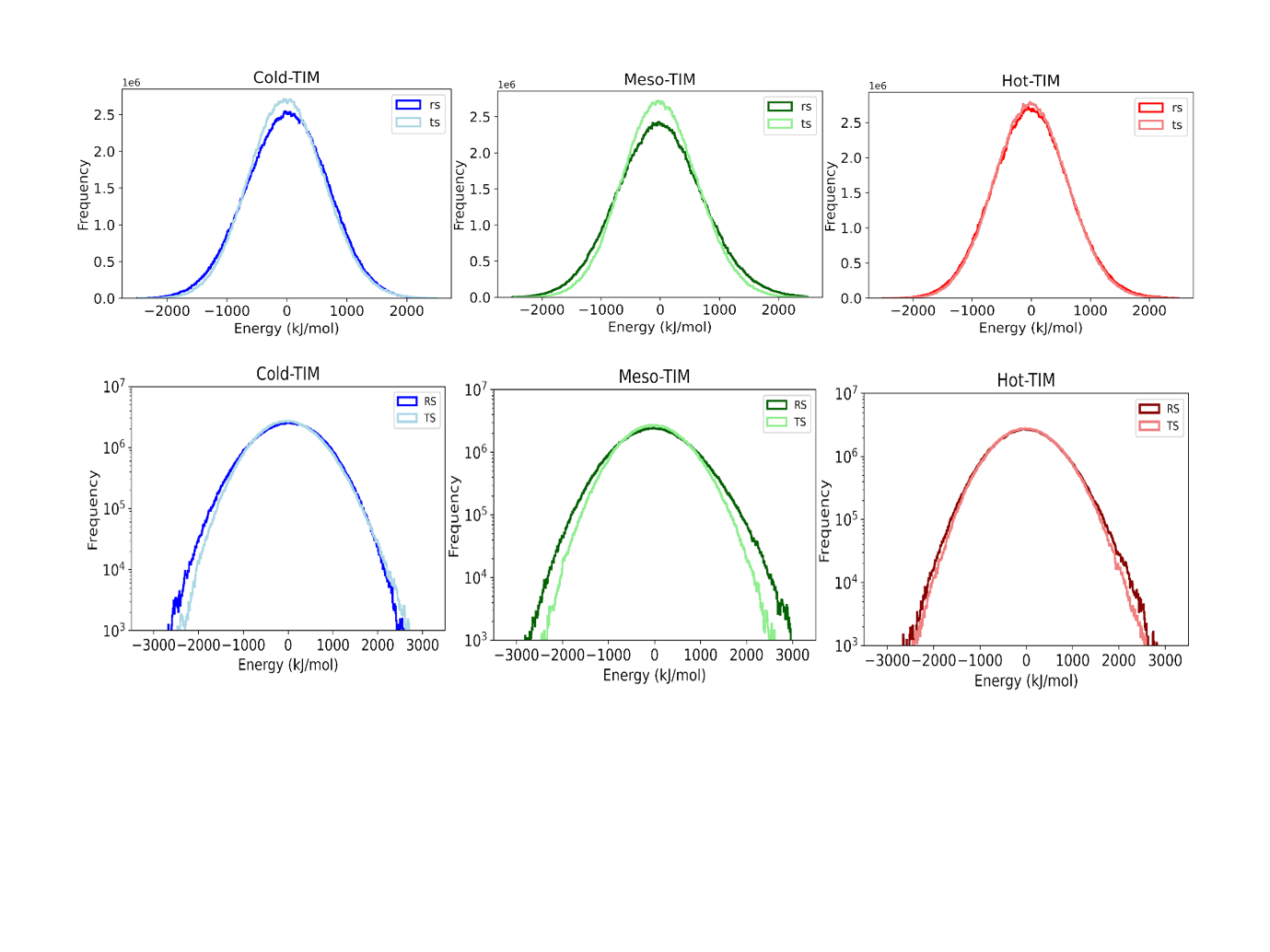

**Figure S9. Potential energy distribution histograms from ten independent 500 ns simulations per state, normalized to a 50 ns moving average.** Dark colours: RS complex; light colours: TS complex. Top row: linear y-axis. Bottom row: logarithmic y-axis to visualize distribution tails. The TS distribution (light) is narrower than RS (dark) in cold-TIM and meso-TIM, yielding negative ${\Delta C}^{\ddagger}$ values.

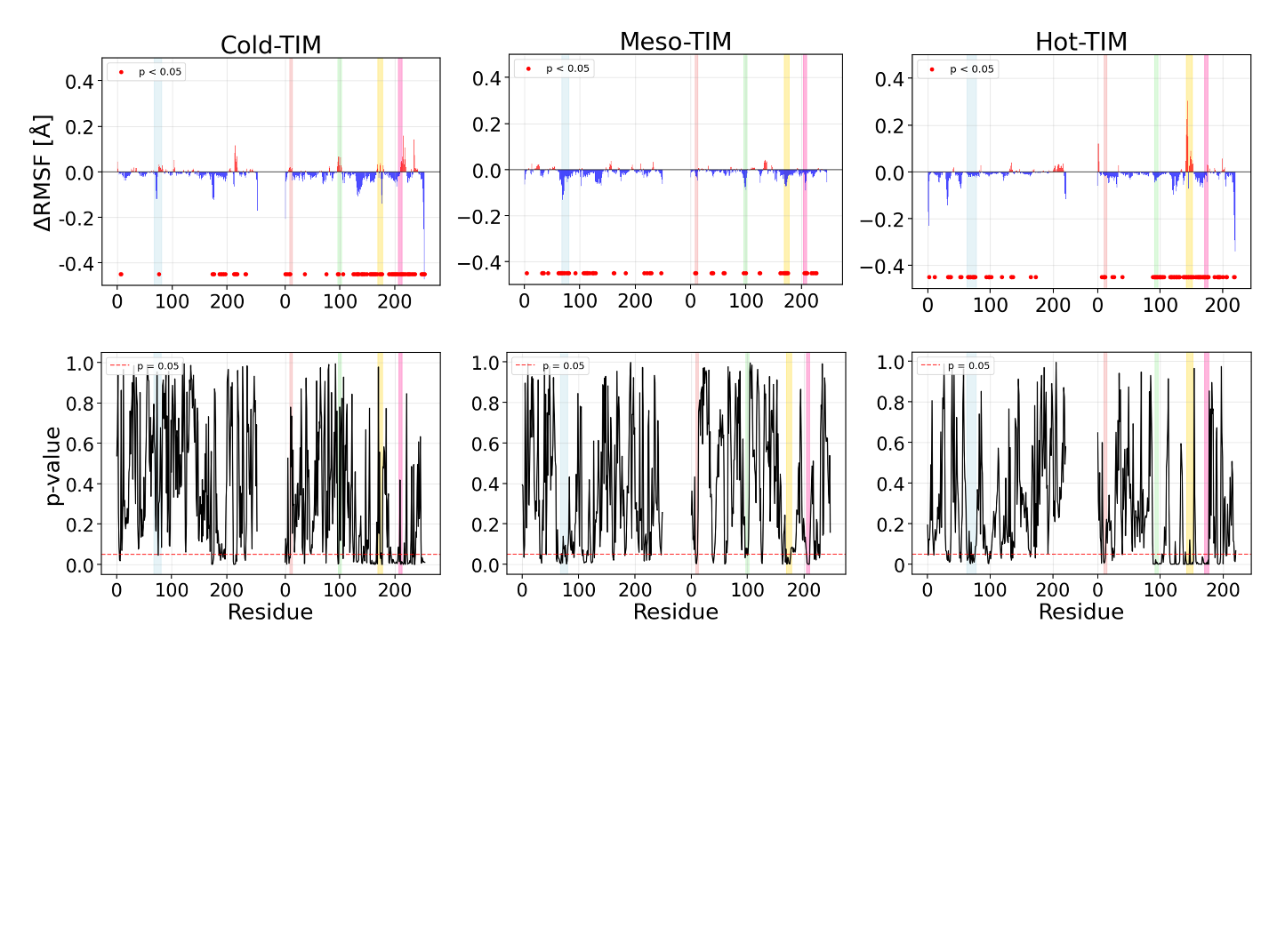

**Figure S10.** **Cα ΔRMSF between RS and TS complexes (for 10ns windows) and their significance (p-value) in each TIM**. Positive (red) values indicate residues that exhibit greater flexibility in TS, while negative (blue) values denote increased flexibility in RS. Significance was determined using a two-sample t-test on Cα RMSF measurements between the two states. A vertical dotted line represents the position of the dimeric interface, while a horizontal dotted line indicates the significance threshold (p < 0.05) as determined by t-test. Residues with statistically significant p-values are denoted by red dots.

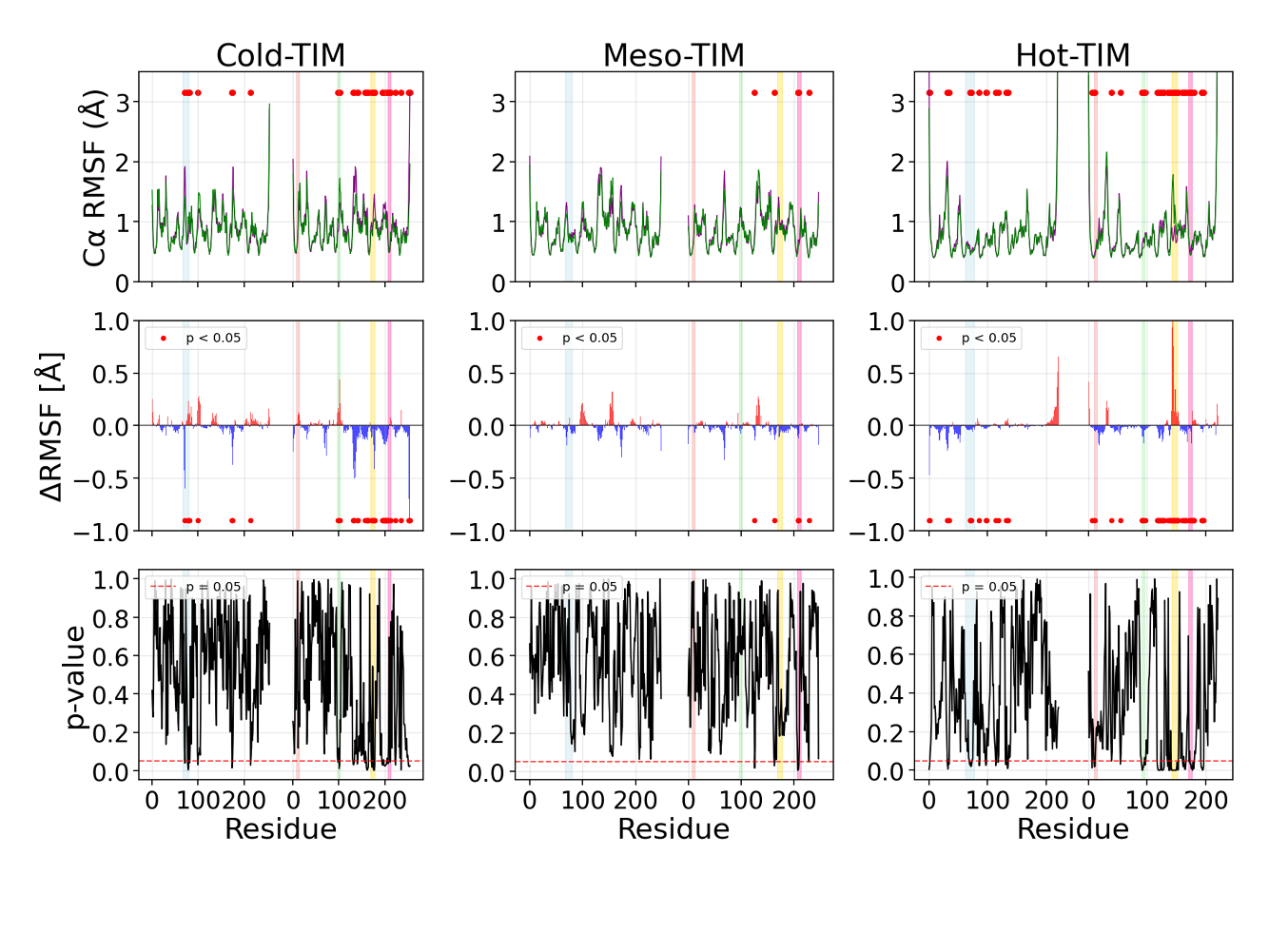

**Figure S11. Cα RMSF (top); ΔRMSF (middle) between RS and significance (p-value, bottom) and (on 450 ns timescale) in each TIM**.**(top)** Average Cα RMSF for RS (purple) and TS (green) states calculated over 450 ns windows. Red dots indicate statistically significant differences (p < 0.05, two-sample t-test). Coloured vertical bars denote motifs 1-5. **(middle)** ΔRMSF (TS - RS) values. Positive (red) indicates increased TS flexibility; negative (blue) indicates increased RS flexibility. Red dots denote significant differences (p < 0.05). (**bottom)** P-values from two-sample t-tests. Horizontal dotted red line indicates significance threshold (p = 0.05).

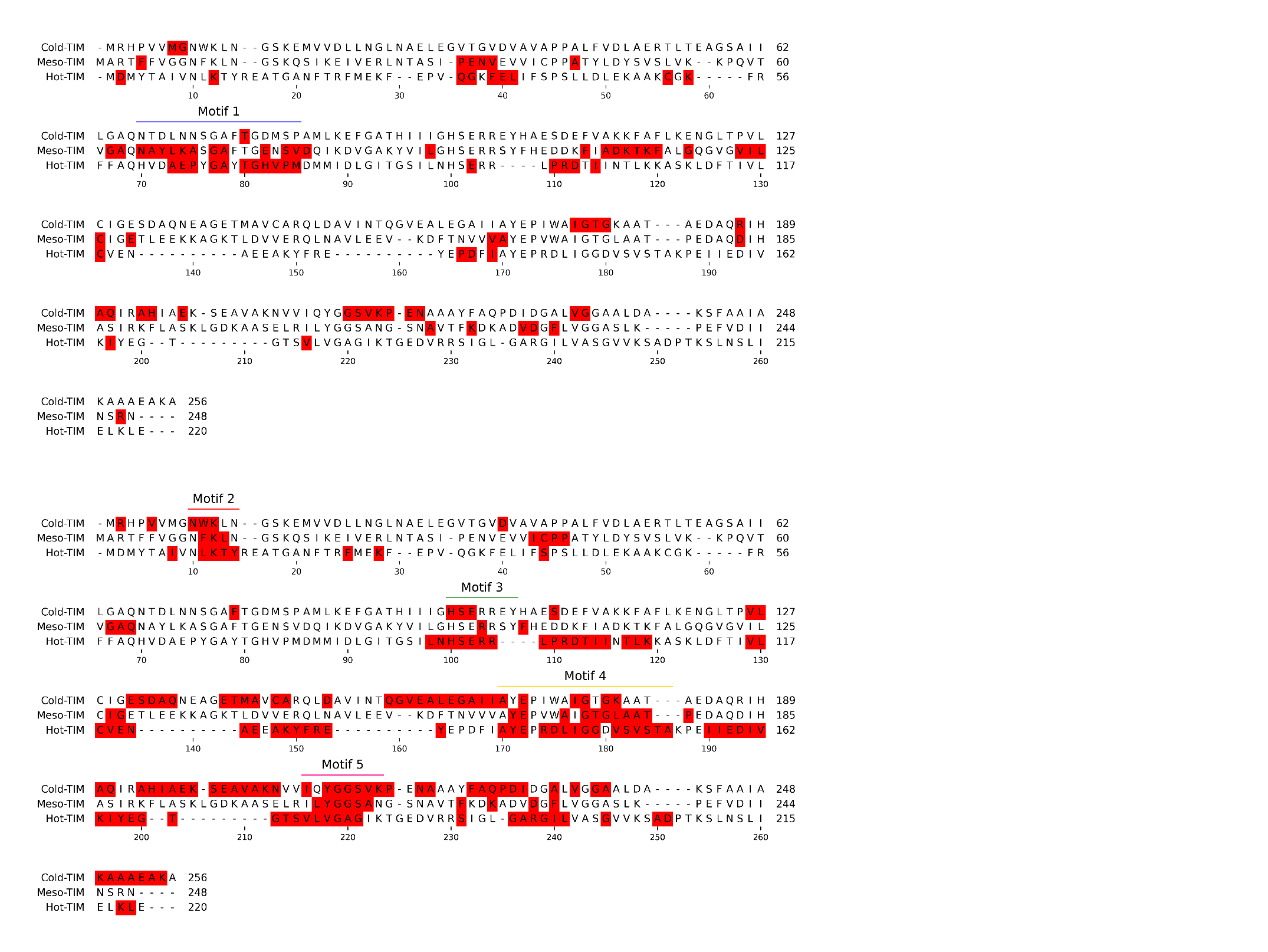

**Figure S12**. **Structure-based sequence alignment of three TIMs.** The alignment was performed using MAFFTv7.0 (17) using default parameters. Residues/motifs highlighted in boxes in the sequence alignment show a significant difference in ΔRMSF as determined by a two-sample t-test. These regions are also highlighted in superimposed TIM structure in **Figure S11**.

|  | Cluster ID | Total Cluster (%) | Cluster fraction RS (%) | Cluster fraction TS (%) |
| --- | --- | --- | --- | --- |
| Hot-TIM | 0 | **94.68** | 97.24 | 92.12 |
|  | 1 | 3.06 | 0.74 | 5.38 |
| Meso-TIM | 0 | **35.88** | 33.4 | 38.35 |
|  | 1 | **19.74** | 31.38 | 8.09 |
|  | 2 | **9.9** | 4.07 | 15.72 |
|  | 3 | **9.38** | 3.84 | 14.92 |
|  | 4 | **8.05** | 8.8 | 7.3 |
|  | 5 | **5.53** | 11.02 | 0.05 |
|  | 6 | 2.56 | 2.6 | 2.52 |
|  | 7 | 2.41 | 0.01 | 4.82 |
|  | 8 | 2.19 | 0.37 | 4.01 |
| Cold-TIM | 0 | **17.15** | 1.65 | 32.65 |
|  | 1 | **13.75** | 27.49 | 0.01 |
|  | 2 | **11.07** | 21.42 | 0.73 |
|  | 3 | **8.21** | 0.01 | 16.4 |
|  | 4 | **5.66** | 11.19 | 0.12 |
|  | 5 | **5.13** | 0.24 | 10.02 |
|  | 6 | **4.93** | 0.25 | 9.61 |
|  | 7 | **4.76** | 0.01 | 9.51 |
|  | 8 | **4.45** | 8.9 | 0 |
|  | 9 | 3.82 | 7.65 | 0 |
|  | 10 | 3.75 | 0.31 | 7.18 |
|  | 11 | 3.53 | 0.02 | 7.04 |
|  | 12 | 3.33 | 6.57 | 0.09 |
|  | 13 | 2.16 | 4.21 | 0.1 |

**Table S4**. **Combined clustering of RS and TS complexes.** A total of 20 simulations (10 for each state) was analysed using the hierarchical agglomerative clustering algorithm with a minimum cluster distance (epsilon) of 1.5 Å for each TIM was used. This analysis identifies 9 clusters for psychrophilic TIM, 6 clusters for mesophile TIM and only one for the thermophile (with overall contribution >4%). Clusters with fractions >4% are shown in bold.

| **Cutoff** | **Cold-TIM** | **Meso-TIM** | **Hot-TIM** |
| --- | --- | --- | --- |
| 0.05 | 62416 (24.0%) | 44042 (18.1%) | 16367 (8.5%) |
| 0.1 | 47475 (18.3%) | 27435 (11.2%) | 3685 (1.9%) |
| 0.2 | 21069 (8.1%) | 6669 (2.7%) | 203(0.1%) |
| 0.3 | 5166 (2.0%) | 1089 (0.5%) | 14 (0.01%) |
| 0.4 | 472 (0.2%) | 190 (0.08%) | 0 |

**Table S5. Quantitative analysis of correlation changes across TIM variants.** Number and percentage of residue pairs showing increased correlation or increased anti-correlation between reactant state (RS) and transition state (TS) complexes for different correlation coefficient thresholds.

| **Cutoff** | **Cold-TIM** | **Meso-TIM** | **Hot-TIM** |
| --- | --- | --- | --- |
| 0.05 | 11246 (4.3%) | 16545 (6.8%) | 10675 (5.5%) |
| 0.1 | 4997 (1.9%) | 7917 (3.2%) | 2560 (1.3%) |
| 0.2 | 546 (0.2%) | 1510 (0.6%) | 288 (0.2%) |
| 0.3 | 59 (0.02%) | 234 (0.1%) | 57 (0.03%) |
| 0.4 | 10 (0%) | 38 (0.02%) | 20 (0.01%) |

**Table S6. Analysis of correlation losses between reactant and transition states.** Number and percentage of residue pairs showing decreased correlation or decreased anti-correlation between between reactant state (RS) and transition state (TS) complexes at various correlation coefficient thresholds.

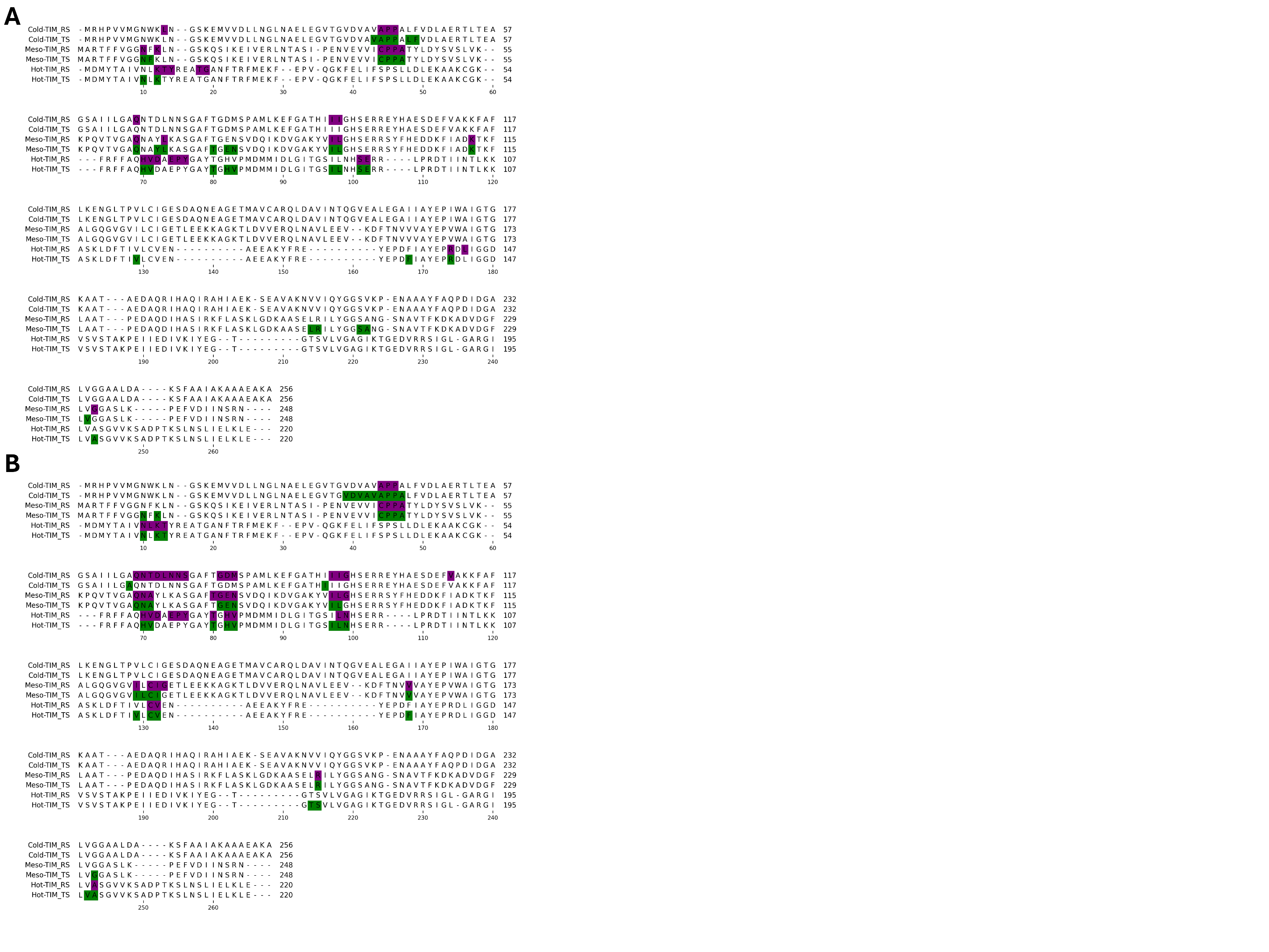

**Figure S13. Shortest Path Map (SPM) network nodes mapped onto multiple sequence alignment of TIM variants.**

Multiple sequence alignment showing residues that participate in dynamical communication networks for cold-TIM meso-TIM and hot-TIM **(A)** Non-reactive monomer: RS network nodes highlighted in purple and TS network nodes highlighted in green. **(B)** Reactive monomer: RS and TS network nodes shown with the same color scheme. Alignment generated using MAFFT v7.0.

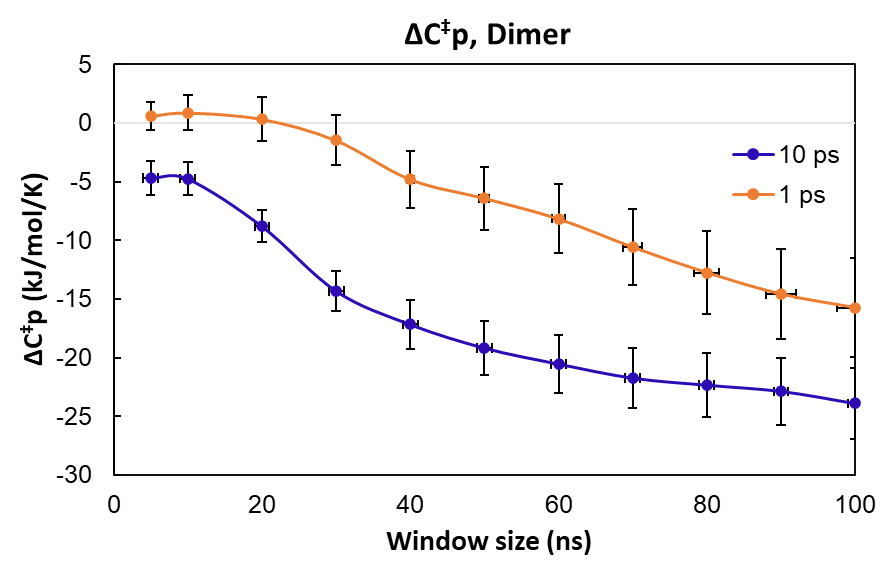

**Figure S14**: **Activation heat capacity values for the psychrophilic TIM dimer, with different values of coupling parameter in the Berendsen thermostat.** Calculations for 10 ps coupling to the thermostat are shown by the blue curve, 1 ps values are shown by the orange curve.

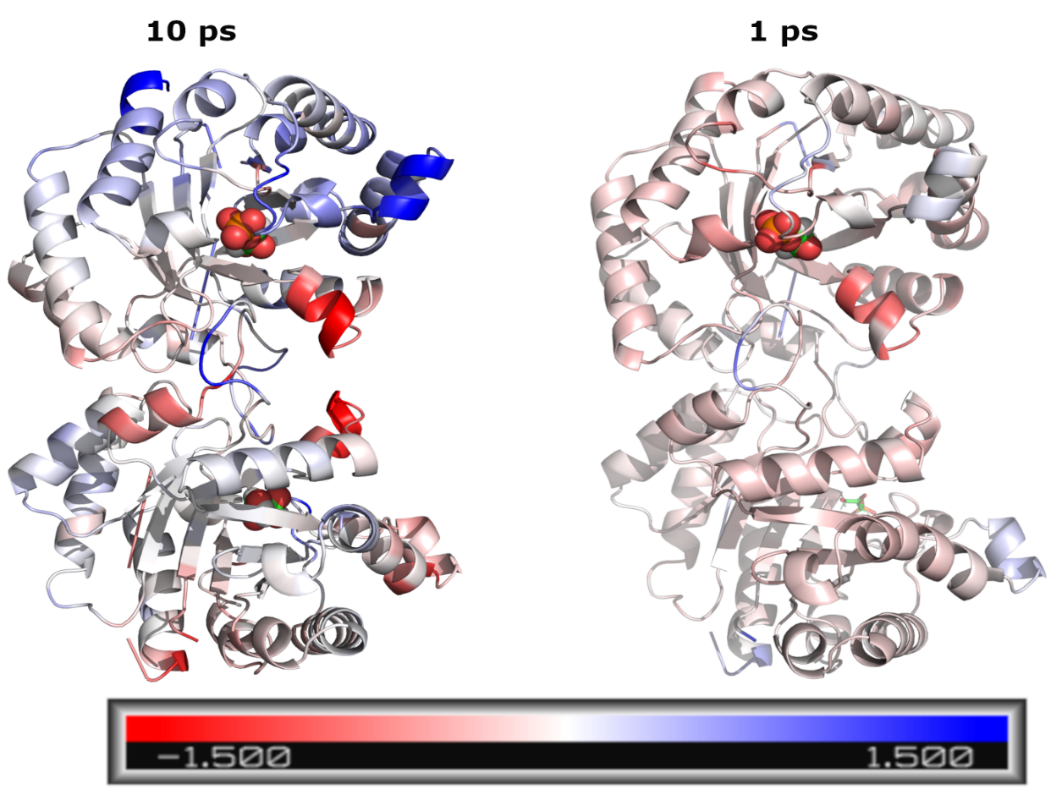

**Figure S15**: **Effect of thermostat coupling parameter values on RMSF**. Mapping of ΔRMSF (RS-TS) values for cold-TIM with 10 ps and 1 ps coupling parameter in the Berendsen thermostat.

Åqvist and van der Ent have criticised the method which we have used for ${\boldsymbol{\Delta}\boldsymbol{C}}^{\ddagger}$calculation (*18*). Additionally, they discussed the problem of ensuring that MD simulations yield correct total energy fluctuations, which are dependent on the thermostat used. Specifically, the use of the Berendsen thermostat, with a 10 ps coupling constant, was questioned, as this may underestimates the energy fluctuations. The deviation becomes larger when the value of the coupling constant, τ, is increased; this is because it weakens coupling to the heat bath. It was concluded that “the fluctuation formula requires careful attention to the thermostat”. To test this, we carried out further simulations using the Berendsen thermostat with a coupling parameter of 1 ps for cold-TIM, and compare the results to the simulations that used a larger coupling parameter of 10 ps.

**Figure S10** shows a direct comparison of${\Delta C}^{\ddagger}$for simulations with different thermostat parameters: blue, 10 ps coupling; orange, 1 ps coupling. The two curves share a similar shape, but the values for the 1 ps simulations are shifted towards less negative values. For example, with a window size of 80 ns, ${\Delta C}^{\ddagger}$is -12.7 kJ/mol/K for 1 ps coupling, compared to –22.3 kJ/mol/K for 10 ps coupling. With the smaller, tighter 1 ps coupling, the temperature of the system is corrected more quickly, which should inhibit fluctuations within the enzyme structure. MMRT theory proposes that fluctuations are greater in the RS than the TS; therefore, it could be reasoned that the tighter coupling constant will have a more evident effect on the RS. Referring to our formula for ${\Delta C}^{\ddagger}$(*eq. 1*), a substantial reduction in RS fluctuations compared to that of the TS will lead to a less negative overall value. On this basis, it can be reasoned that the simulations are following the predictions of MMRT and the fluctuation formula.

Looking at the curve for the 1 ps coupling, the ${\Delta C}^{\ddagger}$values are initially positive, before becoming negative at a window size of 30 ns. For window sizes greater than 30 ns, ${\Delta C}^{\ddagger}$ values continue to become more negative. It could be postulated that the active site loops are opening at a longer timescale, along with increased fluctuations in external regions of the enzyme, both of which would contribute to the more negative ${\Delta C}^{\ddagger}$ values. Further, Åqvist et al. concluded that when using the fluctuation formula, longer simulations are required for good statistics; however, they reported only 30 ns of data for each temperature that was investigated. The results presented here show that a timeframe of 30 ns is too short to represent the dynamics of the enzyme.
